# Cooperative RNA degradation stabilizes intermediate epithelial-mesenchymal states and supports a phenotypic continuum

**DOI:** 10.1101/2022.05.10.491398

**Authors:** Benjamin Nordick, Mary Chae-Yeon Park, Vito Quaranta, Tian Hong

## Abstract

Epithelial-mesenchymal transition (EMT) is a change in cell shape and mobility that occurs during normal development or cancer metastasis. Multiple intermediate EMT states reflecting hybrid epithelial and mesenchymal phenotypes were observed in various physiological and pathological conditions. Previous theoretical models explaining the intermediate EMT states rely on multiple regulatory loops involving transcriptional feedback. These models produce three or four attractors with a given set of rate constants, which is incompatible with experimentally observed non-genetic heterogeneity reflecting a continuum-like EMT spectrum. EMT is regulated by many microRNAs that typically bind transcripts of EMT-related genes via multiple binding sites. It was unclear whether post-transcriptional regulations associated with the microRNA binding sites alone can stabilize intermediate EMT states. Here, we used models describing the post-transcriptional regulations with elementary reaction networks, finding that cooperative RNA degradation via multiple microRNA binding sites can generate four-attractor systems without transcriptional feedback. We identified many specific, experimentally supported instances of network structures predicted to permit intermediate EMT states. Furthermore, transcriptional feedback and the newly identified intermediates-enabling circuits can be combined to produce even more intermediate EMT states in both modular and emergent manners. Finally, multisite-mediated cooperative RNA degradation can increase the distribution of gene expression in the EMT spectrum and support the phenotypic continuum without the need of higher noise. Our work reveals a previously unknown role of cooperative RNA degradation and microRNA in EMT, providing a theoretical framework that can help to bridge the gap between mechanistic models and single-cell experiments.

## Introduction

Epithelial-mesenchymal transition (EMT) is a cell state change required for embryogenesis, postnatal development, and some diseases’ progression including metastasis (1, 2). During EMT, epithelial cells lose their apical-basal polarity and gain the ability to migrate. The transition is not a binary switch: intermediate cellular phenotypes between epithelial (E) and mesenchymal (M) states have been found in development, mammalian cell lines, fibrosis, and tumors (3–7). These intermediate EMT states may possess characteristics of both E and M phenotypes, which can be important for cell- or tissue-level functions such as collective cell migration (8). Recent single-cell transcriptomic studies using individual cell lines provide additional evidence for the existence of multiple intermediate EMT states (5, 9–11). To investigate the mechanistic basis of the intermediate states, theoretical and experimental approaches were used to demonstrate that interconnected transcriptional feedback loops can support intermediate EMT states (6, 12–14), but the existing mechanistic models can produce only one or two intermediate EMT states with a set of biochemical rate constants reflecting one cell type. Contrary to these models, it was recently shown that accurate description of EMT dynamics in a single mammary epithelial cell line requires at least several more intermediate states (15). This finding is consistent with the transcriptomic data showing a continuum-like EMT spectrum which possibly contains many stable intermediate cell states (5, 9–11). These observations indicate that an improved theoretical foundation of the intermediate EMT states is needed to bridge the gap between models and experimental data.

It was shown that EMT involves extensive post-transcriptional regulation by microRNAs (miRNAs): more than one hundred miRNAs were found to be significantly associated with EMT (16, 17). Both epithelial-associated miRNAs and mesenchymal-associated miRNAs have been identified, inhibiting mesenchymal genes (M genes) and epithelial genes (E genes) respectively. Mechanistically, this miRNA-mediated inhibition is through mRNA degradation and translation repression upon miRNA-mRNA binding (18). It was recently shown that multiple epithelial-associated microRNAs can inhibit in a strongly cooperative manner (17), further supporting the roles of miRNAs in EMT. While existing models describing intermediate EMT states often include miRNA regulations, these models require multiple transcriptional feedback loops (6, 12, 13). However, it is unclear whether intermediate EMT states can arise from simpler, more common gene regulatory networks.

In this work, we use mathematical models describing elementary RNA reaction networks to show that cooperative RNA degradation can generate intermediate EMT states in the absence of transcriptional feedback. We use bioinformatic approaches to demonstrate that the topologies of gene regulatory networks allowing multiple intermediate cell states are widespread in the EMT system. Furthermore, transcriptional and post-transcriptional mechanisms can be combined to support larger numbers of intermediate EMT states in both modular and emergent manners. Finally, we use a comprehensive EMT model to show that cooperative RNA degradation can facilitate the formation of a phenotypic continuum. Our work reveals a previously unknown mechanism for intermediate EMT states and provides a new theoretical framework for understanding the perplexing EMT spectrums in development and disease progression.

## Results

### Cooperative RNA degradation generates intermediate EMT states in the absence of transcriptional feedback

Based on recent theories and experiments showing post-transcriptional mechanisms for bistability (19), we hypothesized that intermediate EMT states can arise without transcriptional feedback. To test the hypothesis, we first considered models with mass-action kinetics describing interactions between microRNAs and mRNAs as well as their synthesis and degradation. It was previously proved that with arbitrary positive rate constants, an mRNA with one microRNA binding site (the MMI1 Model) can have only one stable steady state, and that an mRNA with two microRNA binding sites (the MMI2 Model) can have at most two stable steady states (19). In addition, systematic search with biologically relevant parameter sets showed that, like the MMI2 Model, an mRNA with three microRNA binding sites (the MMI3 Model) can have at most two stable steady states (19). We therefore built a model containing an mRNA with four microRNA binding sites (the MMI4 Model). In the first version of the MMI4 Model, the four microRNA binding sites are bound by one microRNA. The two RNAs can form four types of complexes through complementarity-based binding, i.e. 1:*n* (*n* ∈ (1,2,3,4)) complex where *n* represents the number of microRNA molecules in each complex (Figure 1A). The two RNAs are allowed to be degraded independently with distinct rate constants in the complexes. Details of all models can be found in the supplemental information. We randomly sampled parameter values within biologically plausible ranges. Out of 10^7^ sampled parameter sets, we found 1,732 that generated three stable steady states (i.e. three attractors, i.e. tristability, Figure 1B-C) and none that generated four attractors (tetrastability). The parameter sets allowing three attractors contained degradation rate constants that are individually plausible, but collectively unusual. For example, the most extreme degradation rates are often found in unsaturated complexes (example parameter sets of interest can be found in the supplemental information), which would require the mRNA to be strongly stabilized by the binding of additional microRNA molecules. This single-microRNA MMI4 Model is therefore not likely to drive multistability *in vivo*.

**Figure 1.**
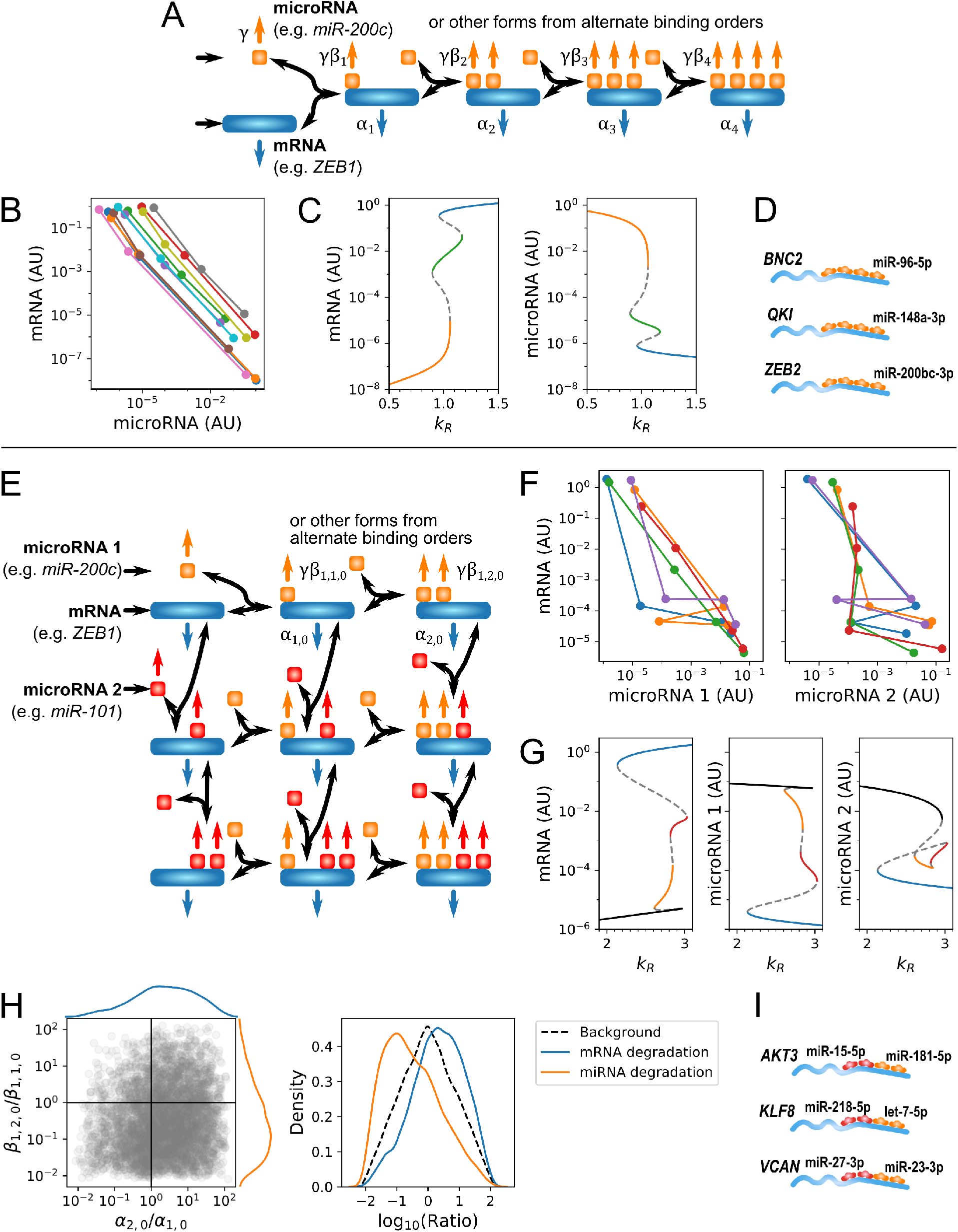
Multistability in the MMI4 Models. (**A**) Reactions in the one-microRNA MMI4 Model, which has four binding sites for the same microRNA on one mRNA. Wide rectangles, mRNA; squares, microRNA; horizontal arrows, transcription; colored arrows, RNA degradation; curved arrows, binding/unbinding. (**B**) Example tristable systems from the one-microRNA MMI4 Model. Points represent attractors in the space of free mRNA vs. free microRNA concentration. Attractors of the same system/parameterization are joined by lines of the same color. AU, arbitrary units. (**C**) Bifurcation diagrams showing the steady states of free mRNA (left) and free microRNA (right) as a function of the mRNA transcription rate *k_R_* from the brown system in B. Each steady state is colored the same in both plots. Dashed lines, unstable steady states. Parameter values in supplemental information. (**D**) Other EMT-related examples of the one-microRNA MMI4 Model. (**E**) Reactions in the two-microRNA MMI4 Model, which has two binding sites for each of two microRNAs on one mRNA. (**F**) Example tetrastable systems from the two-microRNA MMI4 Model in the space of free mRNA vs. free microRNA 1 (left) or free microRNA 2 (right). Order of attractors connected by each line is the same in both subplots. (**G**) Bifurcation diagrams showing the steady states of free mRNA (left), free microRNA 1 (middle), and free microRNA 2 (right) as a function of the mRNA transcription rate from the green system in F. Parameter values in supplemental information. (**H**) Left: Scatterplot of functional cooperativities in mRNA degradation (*α*_2,0_/*α*_1,0_) and microRNA 1 degradation (*β*,_1,2,0_/*β*_1,1,0_) rates due to second microRNA 1 binding in 5,000 3- or 4-attractor systems. Marginal distributions are Gaussian kernel density estimates. Right: Comparison of multistable cooperativity distributions for both microRNAs to distribution of 50,000 randomly sampled parameter sets. (**I**) Other EMT-related examples of the two-microRNA MMI4 Model.

While the MMI4 Model with single microRNA is applicable to EMT (e.g. miR-200c has five putative binding sites on the 3’UTR of *ZEB1* (20), more examples in Figure 1D), allowing the four binding sites to be bound by different microRNAs in the model makes it more flexible to capture a greater number of instances of post-transcriptional EMT circuits with potentially more realistic parameter sets. We therefore considered a modified version of the MMI4 Model which describes two microRNAs each with two binding sites on the mRNA (Figure 1E). This structure indeed allowed the MMI4 Model to produce three- and four-attractor systems with more biologically plausible parameter progressions. These systems contain intermediate EMT states that may be related to the widely observed phenotypes (Figure 1F-G). Almost all parameter sets permitting intermediate EMT states, 4,973 out of 5,000 systems with at least three attractors, involved cooperative degradation of RNAs. The cooperativity is generated by the enhanced degradation rate constant of the mRNA in 1:2 complex with a microRNA compared to the 1:1 complex or by reduced degradation rate constant of the microRNA in 1:2 complex compared to the 1:1 complex (Figure 1H, Figure S1). Cooperative mRNA degradation is observed experimentally (21); functionally cooperative microRNA stabilization might arise through steric blocking of microRNA-degrading factors (22). Three instances of the two-microRNA MMI4 Model are shown in Figure 1I.

EMT is regulated by a large gene regulatory network, in which many genes are targeted by multiple microRNAs (6, 12, 13, 23). Furthermore, the MMI2 Model can generate bistable systems (19). We therefore asked whether connecting two MMI2 modules (the Chained-MMI2 Model, Figure 2A) can enable intermediate EMT states. While this single connection can represent an EMT transcription factor regulating another, this network still does not contain any transcriptional feedback. We found that the Chained-MMI2 Model was able to generate intermediate EMT states in terms of the target gene expression (Figure 2B). We next considered another scenario where one EMT gene is regulated by two EMT transcription factors, each involved in an MMI2 module (the Co-targeting-MMI2 Model, Figure 2C). With this model, intermediate EMT states were observed in terms of the expression of the final target EMT gene (Figure 2D), arising in a combinatorial manner from the multiple upstream bistable systems.

**Figure 2.**
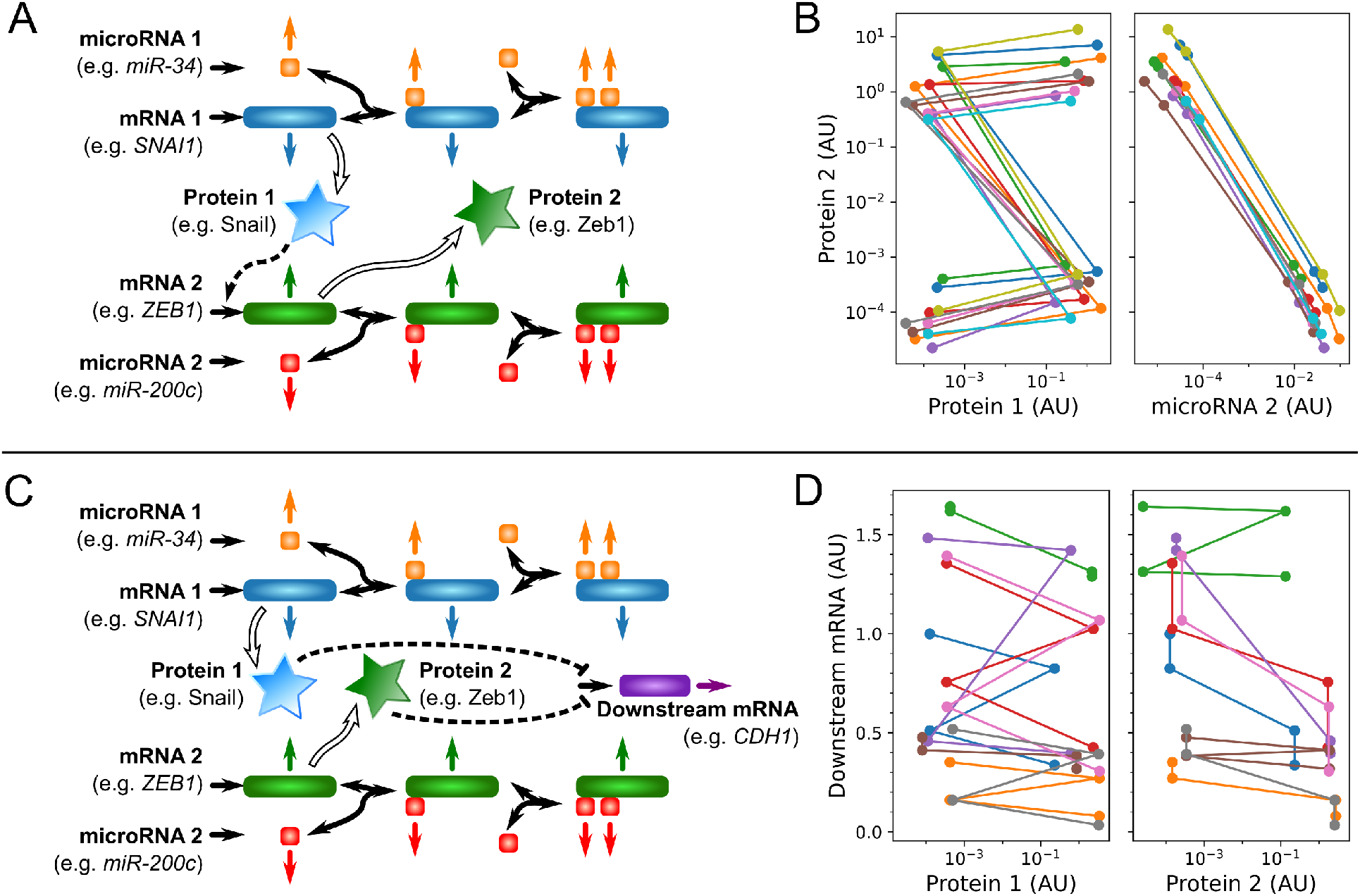
Tetrastability from transcriptionally connected MMI2 targets. (**A**) Reactions in the Chained-MMI2 Model, in which the protein product of one MMI2 target gene transcriptionally regulates another MMI2 target gene. Either microRNA site on each mRNA is allowed to be bound first. Hollow arrows, translation; dashed pointed arrow, transcriptional activation; stars, proteins. (**B**) Example tetrastable systems from the Chained-MMI2 Model in the space of Protein 2 vs. Protein 1 (left) or free microRNA 2 (right). Each system has four different expression levels of Protein 2, which are monotonically anticorrelated to free microRNA 2 levels. (**C**) Reactions in the Co-targeting-MMI2 Model, in which two MMI2 target genes encode proteins that both transcriptionally regulate a third downstream gene. Either microRNA site on each mRNA is allowed to be bound first. Dashed blunt arrow, transcriptional repression. (**D**) Example tetrastable systems from the Co-targeting-MMI2 Model in the space of the downstream mRNA expression vs. Protein 1 (left) or Protein 2 (right). Each system has four different expression levels of the downstream gene.

In summary, we showed that a model with a total of four microRNA binding sites on either one or two EMT genes can generate intermediate EMT states. In each of the four versions of the model, cooperative RNA degradation supports the formation of these intermediate states.

### Network topologies permitting intermediate states driven by post-transcriptional mechanisms are common in EMT regulation

We have shown some specific examples of the intermediates-enabling EMT circuits, but how generally common are these circuits in the EMT system? To address this question, we examined a previously curated list of core EMT genes (24). Among them, 232 were classified as E genes and 191 as M genes. In addition, we used a list of 133 microRNAs, each with experimental evidence supporting its role in EMT (17). With these lists and the TargetScan program for microRNA binding site prediction (20), we first identified 46 EMT genes regulated by a total of four binding sites of one or two EMT microRNAs (Figure 1D/I, Figure S2).

To enumerate the instances of the dual MMI2 Models (Figure 2) that permit intermediate EMT states, we used TRRUST2 and OmniPath, two databases containing experimentally supported transcriptional regulations (25, 26). We first limited the search to *direct* regulations of one EMT gene by another, each involved in an MMI2 module (the Chained-MMI2 Model, Figure 2A), finding 8 instances of this topology. In addition, we found 18 instances of the Co-targeting-MMI2 Model (Figure 2C, Figure S3), again with the assumption of direct regulation. When we changed the assumption to regulation with up to 5 regulatory ‘edges’, we found 171 and 1,312 instances ofthe Chained-MMI2 Model and the Co-targeting-MMI2 Model respectively. Note that these distinct instances are only based on different combinations of EMT genes. Considering combinations of EMT-associated microRNA binding sites would yield much larger numbers of instances.

Overall, our results indicate that the post-transcriptionally driven, intermediates-enabling topologies are common in the EMT system. In fact, 55 of 423 classically defined EMT genes are involved in direct-regulation topologies.

### Modular and emergent synergies between post-transcriptional and transcriptional networks in generating EMT states

It is well known that transcriptional positive feedback loops can generate bistability (27). This type of feedback loop is common in the core EMT regulatory network (6). We next asked whether combining a module consisting of a transcriptional feedback loop (abbreviated as transcriptional module) with a post-transcriptionally driven, intermediates-enabling module (abbreviated as post-transcriptional module) can generate even more intermediate EMT states (28). To test this, we first selected a bistability-enabling parameter set for a representative transcriptional module containing two mutually activating transcription factors (Figure 3A Module 1). We next selected a parameter set for a tetrastability-enabling post-transcriptional module, the Chained-MMI2 Model (Figure 2A, Figure 3A Module 2). Without altering the values of parameters unique to each module, we considered new values for the parameters that conflict, namely the transcription and translation rates of the upstream gene in the Chained-MMI2 Model. We found that there existed values between each pair of original models’ values that allowed the addition of at least one intermediate state to the existing states generated by the post-transcriptional module (Figure 3B). It is remarkable that the addition of intermediate states was achieved without altering the biochemical rate constants within each module, suggesting feasibility of this phenotypic change through evolution.

**Figure 3.**
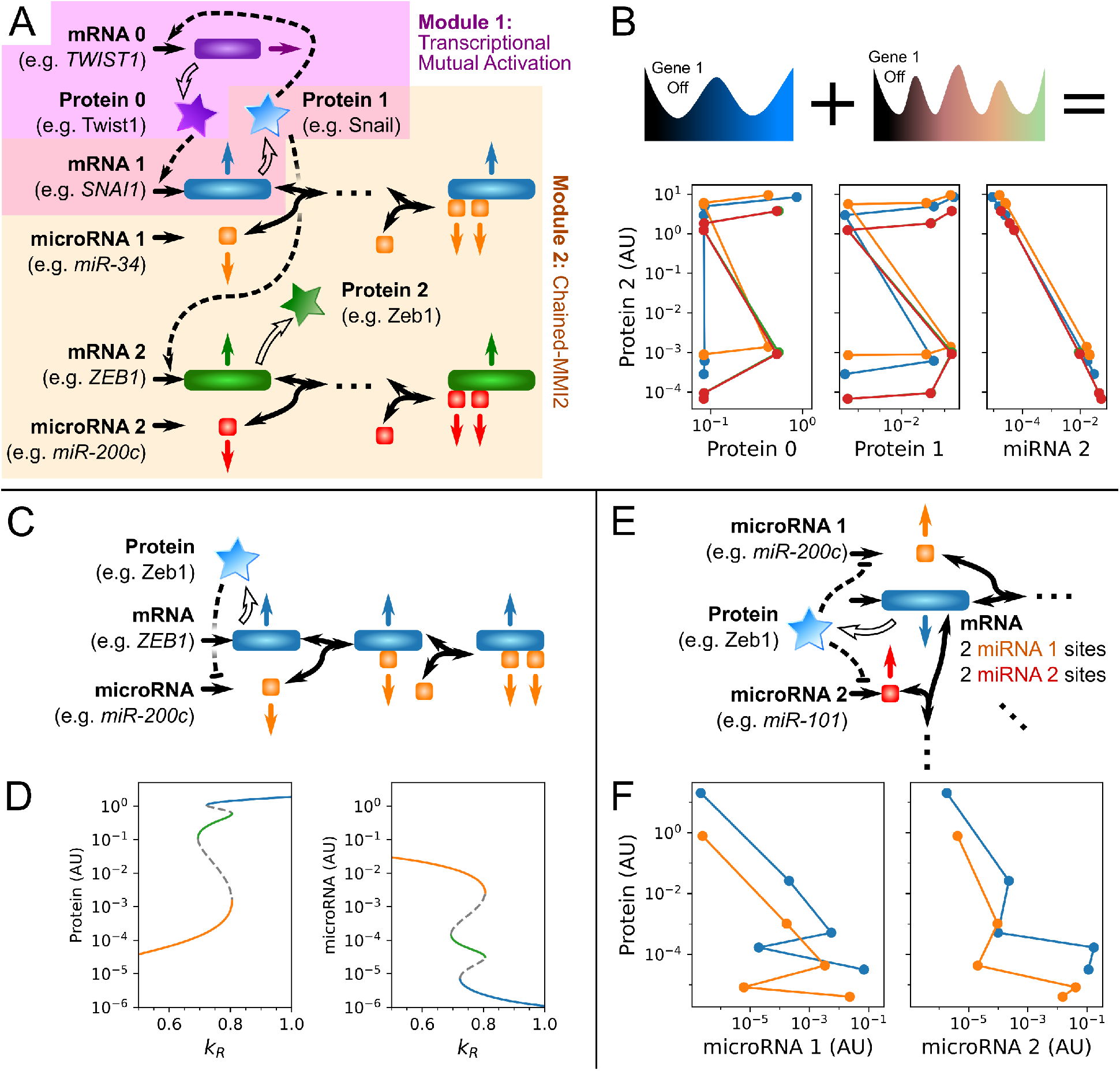
Synergies between transcriptional and post-transcriptional multistability. (**A**) The model resulting from combining a transcriptional mutual activation module (genes 0 and 1, purple shading) with a Chained-MMI2 module (genes 1 and 2, orange shading). (**B**) Example five- and six-attractor systems produced by combining bistable transcriptional mutual activation parameter sets with tetrastable Chained-MMI2 parameter sets. All concentrations are in arbitrary units. (**C**) The model resulting from adding transcriptional repression of the microRNA to the MMI2 Model. (**D**) Bifurcation diagrams showing the steady states of protein (left) and free microRNA (right) levels with respect to the mRNA transcription rate. Tristability can emerge from the addition of the transcriptional repression to MMI2. (**E**) The addition of transcriptional repressions to the two-microRNA MMI4 Model. The mRNA-microRNA complexes are hidden for compactness. (**F**) Example five-attractor systems emerging from the addition of transcriptional repressions to the two-microRNA MMI4 Model.

The modular characteristic of combining transcriptional and post-transcriptional mechanisms for generating multistable systems is applicable to EMT, but there might also exist an “emergent” synergy between the transcriptional and post-transcriptional regulations: a transcriptional module may not be multistable by itself, but when it is combined with a multistable, post-transcriptional module, additional attractor(s) can arise. Indeed, we found that adding a single transcriptional repression of a microRNA to a bistability-enabling MMI2 module involving the same gene gave rise to an intermediate state (Figure 3D). Mechanistically, the emergence of this intermediate cell state was due to the emergent feedback loop between the microRNA and mRNA, consisting of both transcriptional and post-transcriptional regulations (Figure 3C). It was previously shown that this type of hybrid feedback system is common in biology (29). Importantly, the well-known Zeb1-miR200 feedback loop contains this network topology (30). Our results indicate that the previously known tristability-enabling structure containing Zeb1-miR200 feedback loop and an additional transcriptional feedback loop is not the minimal topology for generating an EMT intermediate state (12, 13). Likewise, we found that adding transcriptional repression to the two-microRNA MMI4 Model permits the appearance of a fifth attractor (Figure 3E-F).

In summary, our results suggest that the post-transcriptional and transcriptional networks commonly observed in the EMT system can be combined in both modular and emergent fashions to generate additional intermediate phenotypes.

### A multiple-feedback-mechanism EMT network generates a 7-state EMT continuum

How does the RNA-degradation based mechanism for multistability contribute to the formation of intermediate EMT states in networks larger than simple motifs? To address this question, we selected a basal EMT model published recently: Subbalakshmi et al. showed that a network containing three transcription factors and thirteen transcriptional regulations can generate a three- or four-attractor EMT system with biologically meaningful parameters (31). While there are numerous existing EMT models for tetrastable systems, we chose the Subbalakshmi et al. model because only one microRNA was considered in the model, and it serves as a good basal model for us to test the effect of cooperative RNA degradation by adding more biologically relevant microRNA binding sites sequentially to the system.

As the basal model contains inhibition of *ZEB1* and *SNAI2* by miR-200 without explicit modeling of multiple complexes, we added an additional binding site on *ZEB1* to the model (Figure 4A) and selected a parameter set representing cooperative RNA degradation. As expected, the model with two binding sites of miR-200 produced a system with five attractors (Figure 4B), i.e. an additional intermediate EMT state was generated compared to the previously published results. We next considered another microRNA, miR-101, that regulates *ZEB1* (32) and is repressed by Slug and Snail (23, 33) (Figure 4D). Note that including these additional regulations does not generate any additional transcriptional feedback loops. The inclusion of this microRNA allowed yet two additional intermediate EMT states, resulting in a seven-attractor EMT system (Figure 4E blue).

**Figure 4.**
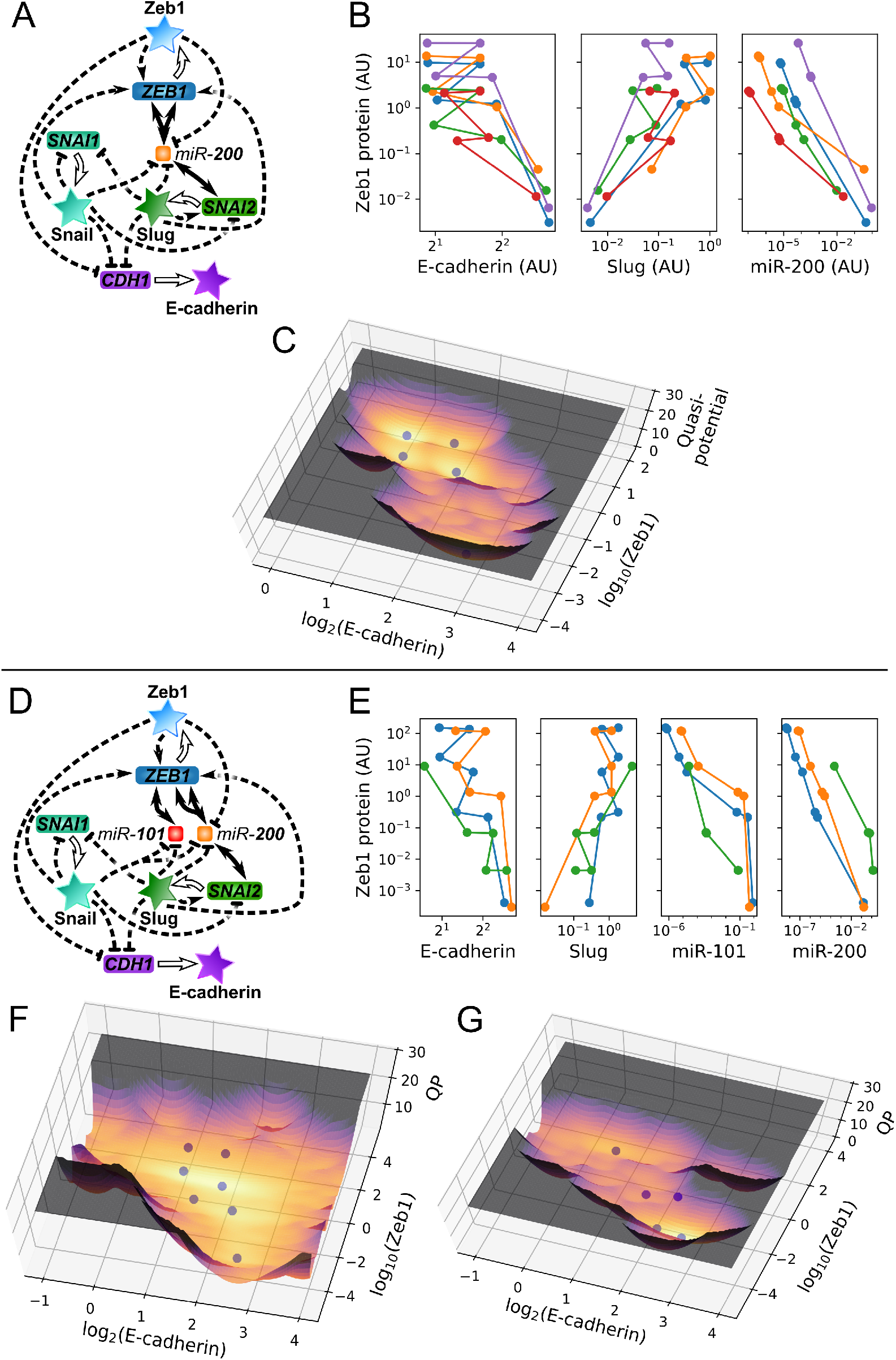
Combining transcriptional and post-transcriptional regulation produces many attractors and a continuum in epithelial-mesenchymal space. (**A**) Structure of the Subbalakshmi et al. model extended with two explicitly modeled *miR-200* binding sites on *ZEB1*. mRNA-microRNA complexes are hidden for compactness. (**B**) Example 5-attractor systems from the model in A shown in the gene expression space of Zeb1 protein vs. E-cadherin protein, Slug protein, or free *miR-200*. (**C**) Quasi-potential diagram showing the stochastic gene expression landscape of the blue system in B under multiplicative noise in the space of E-cadherin protein vs. Zeb1 protein. Deeper regions and brighter color correspond to more likely gene expression states. Spheres, deterministic attractors. (**D**) Structure of the Subbalakshmi et al. model further extended with miR-101 targeting one site on *ZEB1*, transcriptionally repressed by Slug and Snail. (**E**) 7-attractor (blue), 6-attractor (orange), and 5-attractor (green) systems from the model in D. All concentrations are in arbitrary units. (**F**) Quasi-potential (QP) diagram of the 7-attractor blue system in E. (**G**) Quasi-potential diagram of the 5-attractor green system in E.

To test the effect of the cooperative RNA degradation on the EMT system with transcriptional noise, we performed stochastic simulations of the one-microRNA Subbalakshmi et al. network (Figure 4A-C) and the extended model with two microRNAs (Figure 4D-G), applying the same level of multiplicative noise to all RNAs, complexes, and proteins in each model. Based on the steady state distributions of molecules from at least 480 simulations each representing a cell from a population, we constructed the quasi-steady state landscape to visualize the multi-attractor systems under the influence of noise. We found that the cooperative RNA degradation via multiple binding sites not only gave rise to gene expression states near the additional attractors as expected (Figure 4F yellow), but also resulted in broader distributions of gene expression further from the attractors (Figure 4F orange, compare to Figure 4C). While the wider distribution of epithelial and mesenchymal marker genes is not simply an artifact of simulating the more complex model—not every parameter set of the two-microRNA model exhibited such an extreme distribution (Figure 4G)—the new complexes involving the additional microRNA might generally provide more axes in concentration space along which fluctuations, amplified in impact by functional cooperativity in degradation rates, can push the system toward a different state.

In summary, in a larger EMT network containing both transcriptional and post-transcriptional regulations, cooperative RNA degradation via multiple microRNA binding sites gave rise to additional attractors and a broader distribution of gene expression, reflecting the EMT continuum observed in recent single-cell transcriptome data.

## Discussion

Intermediate, or hybrid, EMT phenotypes have been widely observed in several biological contexts. In this work, we have used mathematical models to demonstrate a new mechanism for generating intermediate EMT states based on first principles of gene regulation. This finding can serve as a step towards the reconciliation of the observed EMT continuum with transcriptomic studies and the three or four discrete EMT states captured by previous models. While the observed EMT continuum may be alternatively explained by large subpopulations of cells *en route* to different attractors, recent work using cell-state transition models to explain experimental data showed that models with only a few states cannot describe the time-course EMT data accurately (15). The newly identified post-transcriptional mechanism for generating intermediate states provides a foundation for the additional EMT states necessary to explain expression data at the gene regulation level. A recent study showed that similar post-transcriptional reactions can generate oscillations on slow timescales in addition to multistability (34), which suggests another possibility that the EMT continuum may be supported by a combination of point attractors and cyclic attractors.

Cooperativity of microRNA binding sites has been widely observed (17, 35, 36). In particular, Cursons et al. showed high cooperativity between multiple microRNAs in controlling EMT (17). In this work, we modeled the cooperativity of microRNA binding sites in the form of synergistic mRNA degradation, which was also experimentally observed, and synergistic microRNA stabilization. There are other forms of biologically plausible synergy between binding sites, such as cooperative binding affinities, cooperative translational inhibition, and cooperative inhibition at the network level which involves one microRNA binding to 3’UTRs of multiple mRNAs (e.g. miR-200 in the last model of this study) (21, 37). We expect that some of these mechanisms can be used to support stable intermediate cell phenotypes. Future work is necessary to systematically compare the functions of these molecular mechanisms and to identify their existence in specific biological contexts. Nonetheless, the prevalence of the multi-site microRNA interactions with individual and groups of mRNAs in EMT and other systems suggests that these regulatory networks have nontrivial emergent functions.

In this study, we showed similar and modularizable performances of transcriptional and post-transcriptional mechanisms in generating intermediate EMT states. These two mechanisms are different in their cellular locations. While it may be beneficial for cells to combine both nuclear (transcriptional) and cytosolic (post-transcriptional) machineries to achieve the desired goal of stabilizing intermediate states, the post-transcriptional mechanism may be advantageous in terms of avoiding some sources of noise. This is because transcription is subject to significant noise levels due to the low numbers of DNA coding for the regulatory products, whereas post-transcriptional mechanisms involve large numbers of molecules (38), which reduces intrinsic noise. Therefore, we expect that the proposed RNA-centric mechanism for stabilizing intermediate EMT states can be an efficient strategy for cells to adopt hybrid phenotypes with cytosolic reactions without the need for transcriptional regulatory systems.

The motivation to build our models derived from inconsistencies between existing EMT models that predict a paucity of EMT intermediate states, and experimental single-cell transcriptomic data that have been interpreted to support a wealth of states in a phenotypic continuum (10, 11, 39). Nonetheless, it is plausible that some of the intermediate states should be favored, perhaps in relation with microenvironmental factors such as nutrient availability and cytokines. The models we present here can guide experimentation designed to validate the role of microRNAs to stabilize a constrained number of intermediate cell phenotypes both in physiologic and pathologic systems beyond EMT. For example, in Small Cell Lung Cancer, a phenotypic continuum spanning neuroendocrine (NE) and non-neuroendocrine (non-NE) cell states was recently described based on Archetype Analysis (AA) of experimental data (40, 41). NE to non-NE transitions (NnNT) bear many similarities to EMT, particularly as it pertains increased metastatic properties of non-NE cells, and the similarity of transcriptional signatures (42). Similar to the EMT phenotypic continuum, AA identified a phenotypic continuum for SCLC NE and non-NE subtypes, which gene expression enrichment links to cellular task they are optimized for, such as secretion, proliferation or motility. In this continuum, cells may be specialist at one task, or be suboptimal generalists at one or more tasks. Thus, the intermediate generalist phenotypes arise from task trade-offs, so that they can perform suboptimally to be in tune with the microenvironment that they experience. The continuum then becomes dominated by Pareto optimality. Interrogating the role of microRNAs in NnNT intermediates will provide mechanistic underpinnings for plasticity and high metastatic propensity of SCLC tumors. Our modeling approach is an excellent starting point for streamlining these experiments. In summary, future studies may reveal that post-transcriptional mechanisms are widely used by mammalian cells for generating intermediate states, both for EMT and other differentiation systems. Our mathematical models will aid in designing experiments to test this possibility.

## Methods

### Model construction

Each model is described by a system of ordinary differential equations (ODEs). Dynamics of molecular species are described by two types of function: mass-action kinetics is used to model elementary reactions, i.e. the basal processes of RNA transcription/maturation, constitutive RNA decay, mRNA-microRNA binding and unbinding, regulated decay of each RNA member of each complex, translation, and protein decay; if applicable to the model, Hill functions are used to describe regulated transcription rates as in HiLoop models (43). The models based on the Subbalakshmi et al. network follow the form used there and by Lu et al. (a slightly modified form of Hill functions) (13, 31). In all models, each type of RNA within each type of complex is assigned a regulated degradation factor representing the change in its degradation rate in the complex relative to its free state (34). Binding and unbinding in RNA complexes is always modeled explicitly to ensure that each microRNA molecule acts on only one mRNA at once, though Subbalakshmi et al.’s allowance of incomplete translation repression by microRNA binding (44) is followed.

For simulation, the set of reactions is converted to a system of ODEs, one for each RNA, protein, or complex. A species’ rate of change is the sum of the rates of the reactions that produce or consume it, weighted by the net change in the species’ amount caused by the reaction. As an example, the ODEs for the one-microRNA MMI4 Model are

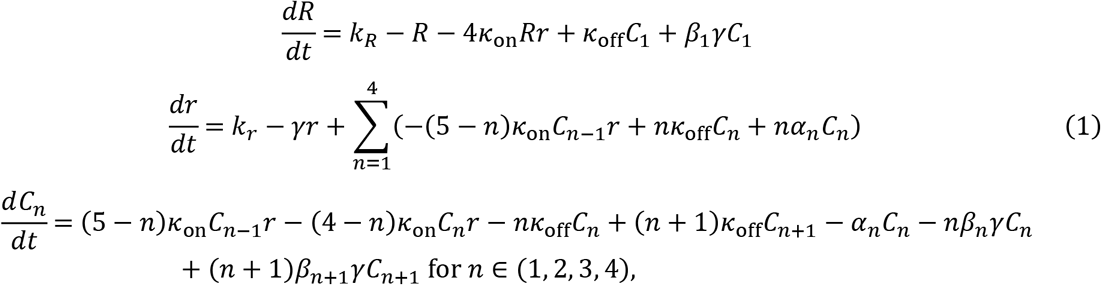

where *R* is the concentration of free mRNA, *r* is the concentration of free microRNA, and *C_n_* is the total concentration of complexes with *n* microRNA molecules bound to an mRNA molecule except *C*_0_ = *R* and *C*_5_ := 0. The regulated mRNA decay reactions, for example, which occur at rate *a_n_C_n_*, destroy one unit of complex *C_n_* but produce *n* units of free microRNA *r. k_R_* and *κ_r_* are transcription rate constants for the two RNAs. *κ*_on_ and *κ*_off_ are association and dissociation rate constants respectively. *γ* is the decay rate constant of free microRNA relative to the free mRNA. *α_n_* and *β_n_* (defined as regulated degradation factors) describe how fast the mRNA and microRNA, respectively, are degraded in the 1:*n* complex relative to their free forms. All variables and parameters have been scaled to dimensionless quantities as described in a recent study (34).

The reactions, rates, and parameters in each model are listed in the supplementary material.

### Parameter sampling for identifying multiple attractors

The likelihood of the single-microRNA MMI4 Model to generate multiple attractors was tested by sampling all regulated degradation factors independently from a log-uniform distribution on [2^-3^,2^4^] following previous work (34, 45), the microRNA association constant from a log-uniform distribution on [10^3^,10^6^], and the microRNA transcription rate from a log-uniform distribution on [2^-4^, 2^1^]. The conditions for the two-microRNA MMI4 Model to produce multiple attractors were investigated by sampling from the same parameter regions independently for each microRNA. Kernel density estimates were computed by SciPy (46) with standard settings. When simply searching for example parameter sets rather than characterizing the parameter space, subregions that provided computational efficiency were selected empirically from plausible regions similar to previous work (34). Parameters of interest for each model are provided in the supplementary material.

To test for multiple attractors, each parameterized system was simulated starting from at least 100 initial conditions which span at least five powers of 5 in a log-uniform manner, drawn from a Sobol quasirandom sequence (47). The initial concentration of each free RNA species was determined by one dimension of the Sobol hypercube, protein initial concentrations were set to their coding mRNA’s initial concentration times the translation rate, and other species’ initial concentrations were zero. Deterministic simulation with Tellurium (48) proceeded for at least 100 time units up to a maximum of 1,000 time units until the system reached a steady state as determined by the 2-norm of the derivatives vector falling below 10^-7^. Simulation endpoint concentration vectors *a* and *b* were considered equivalent attractors if their difference’s 2-norm |*a* – *b*|_2_ was less than 10^-4^ times the number of species *n*, or if |*a* – *b*|_2_ < 0.3 min(|*a*|_2_, |*b*|_2_) < 0.01*n*.

### Numerical bifurcation analysis

Bifurcation diagrams were created with the AUTO2000 plugin for Tellurium 2.2.0 (48), scanning backwards from a high monostable signal value.

### Enumeration of instances of network topologies

The number of potential MMI4 EMT target genes was determined simply by testing each EMT-implicated gene (24) for the presence of four binding sites of EMT-implicated microRNAs (17) using TargetScan (20).

For the topologies that combine transcriptional and posttranscription regulation, the TRRUST2 network (25) and transcriptional subgraph of the OmniPath network (26) as preprocessed for HiLoop (43) were added together except for regulations given opposite signs by the two networks. TargetScan was again used to filter the EMT genes to those matching the MMI2 Model. The existence of a regulatory path from each EMT MMI2 gene to every other, up to the specified path length of 1 or 5, was tested in the combined transcriptional network using NetworkX (49). Each ordered pair with such a directed path was considered an instance of the Chained-MMI2 topology.

The list of EMT MMI2 genes that, directly or indirectly via the combined network, transcriptionally regulates each EMT gene was similarly obtained. If an unordered pair of EMT MMI2 genes regulated the same EMT target gene without common dependencies on any intermediate genes, the partially ordered triple of regulators and target was considered an instance of the Co-targeting-MMI2 topology. The condition of no common intermediates avoids counting subnetworks in which one MMI2 gene regulates the target through the other MMI2 gene, an arrangement which may have different dynamics, and non-minimal subnetworks in which the target is downstream of an already co-targeted gene.

### Stochastic simulation and quasi-potential landscape

We performed stochastic simulations for the modified 3-TF model with various microRNA binding sites by adding an independent multiplicative noise term to each ODE. Divergence arising from negative concentrations was reduced by applying the Zero-Reaction remedy (50). Starting with a population of an equal number of cells at each deterministic attractor, we solved the stochastic ODEs at a noise intensity of 0.2 for 200 time units using DifferentialEquationsjl (51).

To visualize the epithelial-mesenchymal gene expression space in which the stochastic system fluctuates in the long term, the concentrations of Zeb1 and E-cadherin protein were extracted for each simulated cell at intervals of 5 time units starting at time 150. To avoid distortions from temporarily negative concentrations, timepoints with either component less than 10^-7^ were not used. Quasi-potential landscapes (52) were rendered with potential *U*(*x*) = – log*P_S_*(*x*), where *P_S_*(*x*) is the probability density function computed by scikit-learn’s 2-dimensional Gaussian kernel density estimate (53) of bandwidth 0.14 on the base-2 logarithm of E-cadherin protein and the base-10 logarithm of Zeb1 protein, as Zeb1’s expression was more variable under the parameter values tested.

## Author Contributions

Conceptualization: TH. Data curation: MCYP and TH. Investigation: BN and TH. Visualization: BN, MCYP, and TH. Supervision: VQ and TH. Writing—original draft: BN, VQ, and TH.

## Acknowledgements

This work was supported by National Institutes of Health grant R01GM140462 to TH.

## Data Availability

Data sharing is not applicable to this article as no experimental datasets were generated or analyzed during this study. Computer code is available online at https://github.com/BenNordick/MMI4. Additional information is available from the corresponding author upon request.

## Conflict of Interest

VQ is co-founder of Parthenon Therapeutics, Inc., and Duet BioSytems, Inc.. The other authors declare no conflict of interest.

