## Supplementary Materials for "Cooperative RNA degradation stabilizes intermediate epithelial-mesenchymal states and supports a phenotypic continuum"

### General modeling information

For all mRNA-microRNA binding events in all models, the microRNA dissociation rate $\kappa_{\text{off}} = 100$ and each association constant $\text{K} = \kappa_{\text{on}}/\kappa_{\text{off}}$.

$\alpha$ and $\beta$ parameters are regulated degradation factors (RDFs) specifying the degradation rate of an RNA molecule in complex relative to its degradation rate when free.

For mRNAs with multiple binding sites for a type of microRNA, any binding/unbinding order is allowed, but complexes that differ only in the positions of sites bound by their microRNAs are grouped together into one variable, which represents the *total* concentration of all forms of the complex. This is accomplished by multiplying the single-microRNA association rate by the number of remaining free sites and multiplying the dissociation or microRNA degradation rate by the number of occupied sites.

### One-microRNA MMI4

This model contains one mRNA $R = C_0$, one microRNA $r$, and four mRNA-microRNA complexes $C_n$ for the number of bound microRNA molecules $n = 1, 2, 3, 4$.

| **Reactants** | **Products** | **Reaction Rate** | **Description** |
|---|---|---|---|
| - | $R$ | $k_R$ | mRNA transcription |
| $R$ | - | $R$ | mRNA decay |
| - | $r$ | $k_r$ | microRNA transcription/maturation |
| $r$ | - | $\gamma r$ | microRNA decay |
| $C_{n-1} + r$ | $C_n$ | $(5 - n)\kappa_{\text{on}} C_{n-1} r$ | mRNA-microRNA binding |
| $C_n$ | $C_{n-1} + r$ | $n\kappa_{\text{off}} C_n$ | mRNA-microRNA unbinding |
| $C_n$ | $nr$ | $\alpha_n C_n$ | Regulated mRNA degradation |
| $C_n$ | $C_{n-1}$ | $n\beta_n \gamma C_n$ | Regulated microRNA degradation |

| **Parameter** | **Description** | **Figure 1C** |
|---|---|---|
| $\alpha_1$ | RDF for mRNA in the 1:1 complex | 0.310 |
| $\alpha_2$ | RDF for mRNA in the 1:2 complex | 6.305 |
| $\alpha_3$ | RDF for mRNA in the 1:3 complex | 0.716 |
| $\alpha_4$ | RDF for mRNA in the 1:4 complex | 5.744 |
| $\beta_1$ | RDF for microRNA in the 1:1 complex | 3.131 |
| $\beta_2$ | RDF for microRNA in the 1:2 complex | 0.861 |
| $\beta_3$ | RDF for microRNA in the 1:3 complex | 1.255 |
| $\beta_4$ | RDF for microRNA in the 1:4 complex | 1.406 |
| $\gamma$ | Basal microRNA degradation rate | 1 |
| $k_r$ | microRNA transcription rate | 1.042 |
| $k_R$ | mRNA transcription rate | Scanned |
| K | mRNA/microRNA association constant | 270321 |

#### Two-microRNA MMI4

This model contains one mRNA $R = C_{0,0}$, two microRNAs $r_i$ for $i = 1, 2$, and eight mRNA-microRNA complexes $C_{m,n}$ corresponding to all nonzero combinations of zero, one, or two of microRNAs 1 and 2 bound to the mRNA. For example, $C_{2,1}$ represents an mRNA molecule bound by two molecules of microRNA 1 and one molecule of microRNA 2.

| **Reactants** | **Products** | **Reaction Rate** | **Description** |
|---|---|---|---|
| - | $R$ | $k_R$ | mRNA transcription |
| $R$ | - | $R$ | mRNA decay |
| - | $r_i$ | $k_{r,i}$ | microRNA transcription/maturation |
| $r_i$ | - | $\gamma r_i$ | microRNA decay |
| $C_{m-1,n} + r_1$ | $C_{m,n}$ | $(3-m)\kappa_{\text{on},1} C_{m-1,n} r_1$ | mRNA-microRNA 1 binding |
| $C_{m,n}$ | $C_{m-1,n} + r_1$ | $m\kappa_{\text{off},1} C_{m,n}$ | mRNA-microRNA 1 unbinding |
| $C_{m,n-1} + r_2$ | $C_{m,n}$ | $(3-n)\kappa_{\text{on},2} C_{m,n-1} r_2$ | mRNA-microRNA 2 binding |
| $C_{m,n}$ | $C_{m,n-1} + r_2$ | $n\kappa_{\text{off},2} C_{m,n}$ | mRNA-microRNA 2 unbinding |
| $C_{m,n}$ | $mr_1 + nr_2$ | $\alpha_{m,n} C_{m,n}$ | Regulated mRNA degradation |
| $C_{m,n}$ | $C_{m-1,n}$ | $m\beta_{1,m,n}\gamma C_{m,n}$ | Regulated microRNA 1 degradation |
| $C_{m,n}$ | $C_{m,n-1}$ | $n\beta_{2,m,n}\gamma C_{m,n}$ | Regulated microRNA 2 degradation |

| Parameter | Description | Fig. 1F Purple | Fig. 1G |
|---|---|---|---|
| $\alpha_{0,1}$ | RDF for mRNA in the 1:0:1 complex (one miRNA 2) | 2.880 | 2.017 |
| $\alpha_{0,2}$ | RDF for mRNA in the 1:0:2 complex | 12.440 | 4.630 |
| $\alpha_{1,0}$ | RDF for mRNA in the 1:1:0 complex | 3.426 | 1.085 |
| $\alpha_{1,1}$ | RDF for mRNA in the 1:1:1 complex | 3.675 | 7.417 |
| $\alpha_{1,2}$ | RDF for mRNA in the 1:1:2 complex | 12.806 | 8.674 |
| $\alpha_{2,0}$ | RDF for mRNA in the 1:2:0 complex | 5.710 | 1.227 |
| $\alpha_{2,1}$ | RDF for mRNA in the 1:2:1 complex | 15.128 | 4.088 |
| $\alpha_{2,2}$ | RDF for mRNA in the 1:2:2 complex | 24.128 | 5.488 |
| $\beta_{2,0,1}$ | RDF for miRNA 2 in the 1:0:1 complex | 0.797 | 0.485 |
| $\beta_{2,0,2}$ | RDF for miRNA 2 in the 1:0:2 complex | 0.180 | 0.135 |
| $\beta_{1,1,0}$ | RDF for miRNA 1 in the 1:1:0 complex | 0.300 | 0.432 |
| $\beta_{1,1,1}$ | RDF for miRNA 1 in the 1:1:1 complex | 0.526 | 0.742 |
| $\beta_{2,1,1}$ | RDF for miRNA 2 in the 1:1:1 complex | 0.772 | 0.303 |
| $\beta_{1,1,2}$ | RDF for miRNA 1 in the 1:1:2 complex | 0.556 | 0.285 |
| $\beta_{2,1,2}$ | RDF for miRNA 2 in the 1:1:2 complex | 0.115 | 0.240 |
| $\beta_{1,2,0}$ | RDF for miRNA 1 in the 1:2:0 complex | 0.045 | 0.072 |
| $\beta_{1,2,1}$ | RDF for miRNA 1 in the 1:2:1 complex | 0.089 | 0.052 |
| $\beta_{2,2,1}$ | RDF for miRNA 2 in the 1:2:1 complex | 1.000 | 0.353 |
| $\beta_{1,2,2}$ | RDF for miRNA 1 in the 1:2:2 complex | 0.083 | 0.068 |
| $\beta_{2,2,2}$ | RDF for miRNA 2 in the 1:2:2 complex | 0.118 | 0.154 |
| $\gamma$ | Basal microRNA degradation rate | 1 | 1 |
| $k_{r,1}$ | microRNA 1 transcription rate | 0.051 | 0.133 |
| $k_{r,2}$ | microRNA 2 transcription rate | 0.068 | 0.177 |
| $k_R$ | mRNA transcription rate | 2.511 | Scanned |
| $K_1$ | Association constant between mRNA and miRNA 1 | 5319 | 49180 |
| $K_2$ | Association constant between mRNA and miRNA 2 | 3989 | 3852 |

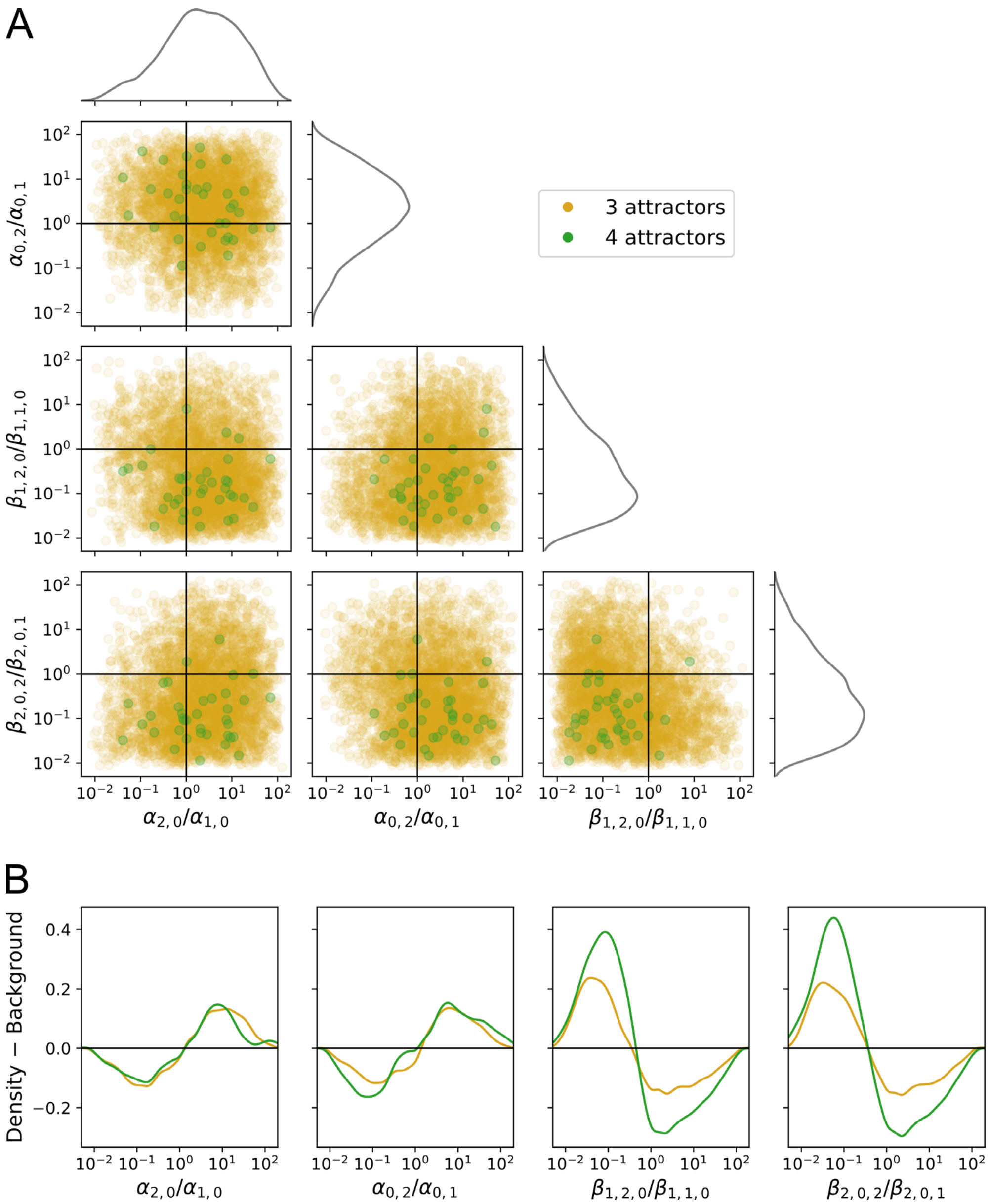

**Figure S1. Functional cooperativity supports multistability in the two-microRNA MMI4 Model.** (**A**) Scatterplots of functional cooperativity in mRNA degradation due to second microRNA 1 binding ($\alpha_{2,0}/\alpha_{1,0}$), mRNA degradation due to second microRNA 2 binding ($\alpha_{0,2}/\alpha_{0,1}$), microRNA degradation due to second microRNA 1 binding ($\beta_{1,2,0}/\beta_{1,1,0}$), and microRNA degradation due to second microRNA 2 binding ($\beta_{2,0,2}/\beta_{2,0,1}$) in 3- or 4-attractor systems found by independently sampling all RDFs from a log-uniform distribution on $[2^{-3}, 2^{4}]$. Marginal distributions are Gaussian kernel density estimates computed from both 3- and 4-attractor systems. (**B**) Differences in kernel density estimate between observed distribution of functional cooperativities for 3- or 4-attractor systems and the distribution obtained by random RDF sampling without testing multistability.

#### Chained-MMI2

This model contains two mRNAs $R_i = C_{i,0}$ for $i = 1,2$, their protein products $X_i$, two microRNAs $r_i$ that each target their mRNA $i$ with two binding sites, and mRNA-microRNA complexes $C_{i,n}$ for the number of bound microRNA molecules $n = 1,2$. Protein 1 activates the transcription of gene 2, with transcriptional activation modeled by a Hill function [1].

| Reactants | Products | Reaction Rate | Description |
|---|---|---|---|
| - | $R_1$ | $k_{R,1}$ | mRNA 1 transcription |
| - | $R_2$ | $k_{R,2}\left(f_{2,C} + \frac{f_{2,1}(X_1/K_{2,1})^{\mathrm{n}_{2,1}}}{1+(X_1/K_{2,1})^{\mathrm{n}_{2,1}}}\right)$ | Regulated mRNA 2 transcription |
| $R_i$ | - | $R_i$ | mRNA decay |
| - | $X_i$ | $l_i R_i$ | Translation |
| $X_i$ | - | $X_i$ | Protein decay |
| - | $r_i$ | $k_{r,i}$ | microRNA transcription/maturation |
| $r_i$ | - | $\gamma r_i$ | microRNA decay |
| $C_{i,n-1} + r_i$ | $C_{i,n}$ | $(3-n)\kappa_{\mathrm{on},i} C_{i,n-1} r_i$ | mRNA-microRNA binding |
| $C_{i,n}$ | $C_{i,n-1} + r_i$ | $n\kappa_{\mathrm{off},i} C_{i,n}$ | mRNA-microRNA unbinding |
| $C_{i,n}$ | $n r_i$ | $\alpha_{i,n} C_{i,n}$ | Regulated mRNA degradation |
| $C_{i,n}$ | $C_{i,n-1}$ | $n\beta_{i,n}\gamma C_{i,n}$ | Regulated microRNA degradation |

| Parameter | Description | Fig. 2B Olive |
|---|---|---|
| $\alpha_{1,1}$ | RDF for mRNA 1 in the 1:1 complex with miRNA 1 | 6.064 |
| $\alpha_{1,2}$ | RDF for mRNA 1 in the 1:2 complex | 18.089 |
| $\alpha_{2,1}$ | RDF for mRNA 2 in the 1:1 complex with miRNA 2 | 1.363 |
| $\alpha_{2,2}$ | RDF for mRNA 2 in the 1:2 complex | 5.334 |
| $\beta_{1,1}$ | RDF for microRNA 1 in the 1:1 complex with mRNA 1 | 0.372 |
| $\beta_{1,2}$ | RDF for microRNA 1 in the 1:2 complex | 0.216 |
| $\beta_{2,1}$ | RDF for microRNA 2 in the 1:1 complex with mRNA 2 | 0.399 |
| $\beta_{2,2}$ | RDF for microRNA 2 in the 1:2 complex | 0.106 |
| $\gamma$ | Basal microRNA degradation rate | 1 |
| $f_{2,C}$ | Proportion of mRNA 2 transcription rate allowed constitutively | 0.545 |
| $f_{2,1}$ | Proportion of mRNA 2 transcription rate regulated by protein 1 | 0.455 |
| $k_{r,1}$ | microRNA 1 transcription rate | 0.043 |
| $k_{r,2}$ | microRNA 2 transcription rate | 0.166 |
| $k_{R,1}$ | mRNA 1 transcription rate | 1.021 |
| $k_{R,2}$ | mRNA 2 maximum transcription rate | 3.123 |
| $\mathrm{K}_1$ | Association constant between mRNA 1 and miRNA 1 | 5000 |
| $\mathrm{K}_2$ | Association constant between mRNA 2 and miRNA 2 | 5000 |
| $K_{2,1}$ | Threshold of the regulation of mRNA 2 by protein 1 | 0.018 |
| $l_1$ | Translation rate for mRNA 1 | 3.552 |
| $l_2$ | Translation rate for mRNA 2 | 5.607 |
| $\mathrm{n}_{2,1}$ | Cooperativity of the regulation of mRNA 2 by protein 1 | 1 |

#### Co-targeting-MMI2

This model contains two microRNA-targeted regulator-encoding mRNAs $R_i = C_{i,0}$ for $i = 1, 2$, their protein products $X_i$, two microRNAs $r_i$ that each target mRNA $i$ with two binding sites, again mRNA-microRNA complexes $C_{i,n}$ for $n = 1, 2$, and a downstream gene's mRNA $R_d$. The downstream gene is subject to transcriptional repression by both regulators, modeled as additively combined Hill functions.

All protein and microRNA-related reactions are as in the Chained-MMI2 Model. Only mRNA dynamics are changed:

| **Reactants** | **Products** | **Reaction Rate** | **Description** |
|---|---|---|---|
| - | $R_i$ | $k_{R,i}$ | Regulators' mRNA transcription |
| - | $R_d$ | $k_{Rd}\left(f_{d,C} + \sum_i \frac{f_{d,i}}{1+(X_i/K_{d,i})^{\mathrm{n}_{d,i}}}\right)$ | Regulated downstream mRNA transcription |
| $R_i$ | - | $R_i$ | Regulators' mRNA decay |
| $R_d$ | - | $R_d$ | Downstream mRNA decay |
| Protein and microRNA-related reactions shared with the Chained-MMI2 Model | | | |

Interpreting gene 1 as *SNAI1*, gene 2 as *ZEB1*, and the downstream gene as *CDH1*, transcription control proportions were estimated based on molecular biology studies [2, 3].

| **Parameter** | **Description** | **Fig. 2D Blue** |
|---|---|---|
| $\alpha_{1,1}$ | RDF for mRNA 1 in the 1:1 complex with miRNA 1 | 5 |
| $\alpha_{1,2}$ | RDF for mRNA 1 in the 1:2 complex | 10 |
| $\alpha_{2,1}$ | RDF for mRNA 2 in the 1:1 complex with miRNA 2 | 5 |
| $\alpha_{2,2}$ | RDF for mRNA 2 in the 1:2 complex | 10 |
| $\beta_{1,1}$ | RDF for microRNA 1 in the 1:1 complex with mRNA 1 | 0.65 |
| $\beta_{1,2}$ | RDF for microRNA 1 in the 1:2 complex | 0.40 |
| $\beta_{2,1}$ | RDF for microRNA 2 in the 1:1 complex with mRNA 2 | 0.65 |
| $\beta_{2,2}$ | RDF for microRNA 2 in the 1:2 complex | 0.40 |
| $\gamma$ | Basal microRNA degradation rate | 1 |
| $f_{d,C}$ | Proportion of downstream transcription rate allowed constitutively | 0.05 |
| $f_{d,1}$ | Proportion of downstream transcription rate regulated by protein 1 | 0.70 |
| $f_{d,2}$ | Proportion of downstream transcription rate regulated by protein 2 | 0.25 |
| $k_{r,1}$ | microRNA 1 transcription rate | 0.1 |
| $k_{r,2}$ | microRNA 2 transcription rate | 0.1 |
| $k_{Rd}$ | Downstream mRNA maximum transcription rate | 1 |
| $k_{R,1}$ | mRNA 1 transcription rate | 1 |
| $k_{R,2}$ | mRNA 2 transcription rate | 1 |
| $\mathrm{K}_1$ | Association constant between mRNA 1 and miRNA 1 | 5000 |
| $\mathrm{K}_2$ | Association constant between mRNA 2 and miRNA 2 | 5000 |
| $K_{d.1}$ | Threshold of the regulation of downstream mRNA by protein 1 | 0.1 |
| $K_{d.2}$ | Threshold of the regulation of downstream mRNA by protein 2 | 0.1 |
| $l_1$ | Translation rate for mRNA 1 | 2 |
| $l_2$ | Translation rate for mRNA 2 | 2 |
| $\mathrm{n}_{d,1}$ | Cooperativity of the regulation of downstream mRNA by protein 1 | 1 |
| $\mathrm{n}_{d,2}$ | Cooperativity of the regulation of downstream mRNA by protein 2 | 1 |

### Instances of RNA-degradation-centric EMT circuits

Seven EMT genes involved in the one-microRNA MMI4 Model structure were identified with TargetScan to predict microRNA binding sites for human genes, a list of EMT-related microRNAs, and a list of EMT-related genes curated earlier (Figure S2A). With a similar approach, we identified 45 EMT genes predicted to be involved in the two-microRNA MMI4 Model structure (Figure S2B).

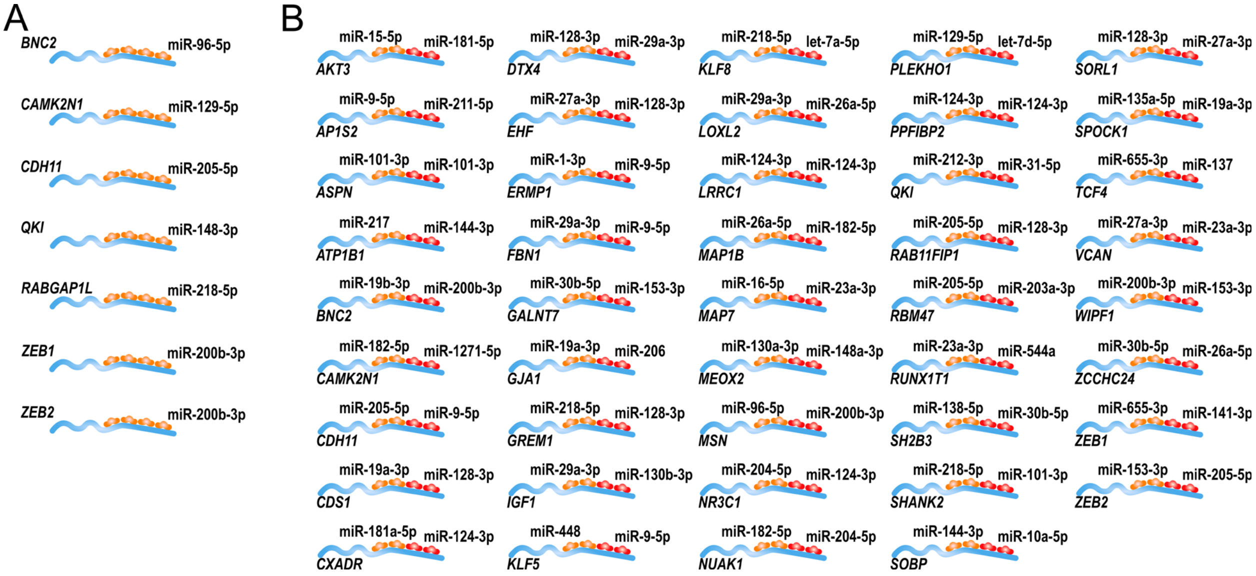

**Figure S2. Instances of the One-microRNA and the Two-microRNA MMI4 Models.** (**A**) Instances of the one-microRNA MMI4 Model. (**B**) Instances of the two-microRNA MMI4 Model. Lists of genes are exhaustive based on our search criteria (see Methods), but only selected examples of the microRNA binding sites are shown.

To estimate the instances of the Chained-MMI2 Model, we looked for direct regulations between EMT genes whose predicted binding sites for EMT microRNAs satisfy the MMI2 Model structure (see Methods). The 8 instances of this model are illustrated in Figure S3A. Similarly, we identified EMT genes involved in the Co-targeting-MMI2 Model structure, and the 18 instances of this model are shown in Figure S3B.

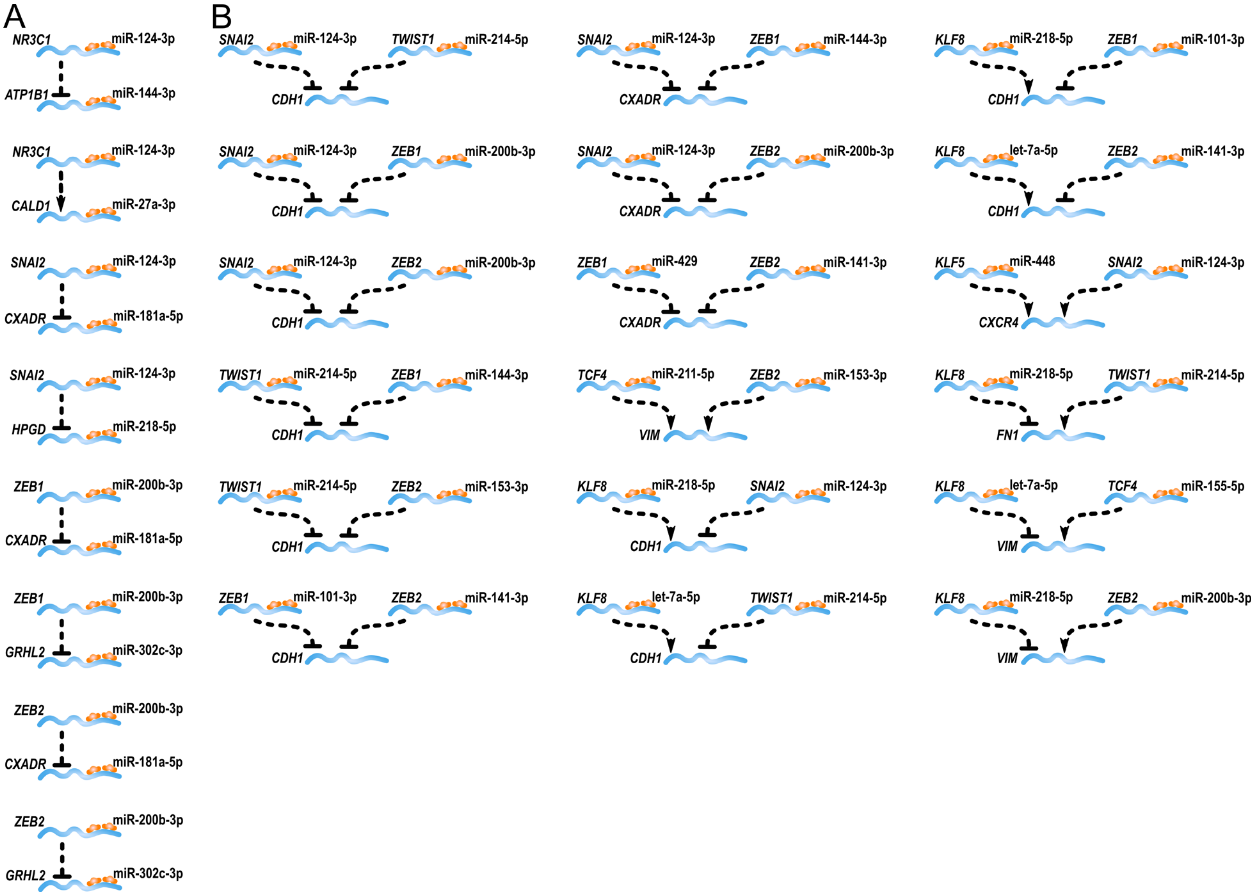

**Figure S3. Instances of the Chained- and Co-targeting-MMI2 Models.** (**A**) Instances of the Chained-MMI2 Model. (**B**) Instances of the Co-targeting-MMI2 Model. Lists of genes are exhaustive based on our search criteria (see Methods), but only selected examples of the microRNA binding sites are shown.

#### Chained-MMI2 with transcriptional positive feedback loop

The transcriptional module consists of two mRNAs $R_j$ for $j = 0, 1$ and their protein products $X_j$:

| Reactants | Products | Reaction Rate | Description |
|---|---|---|---|
| - | $R_j$ | $k_{R,j}\left(f_{j,C} + \frac{f_{j,1-j}\left(X_{1-j}/K_{j,1-j}\right)^{\mathrm{n}_{j,1-j}}}{1+\left(X_{1-j}/K_{j,1-j}\right)^{\mathrm{n}_{j,1-j}}}\right)$ | Regulated mRNA transcription |
| $R_j$ | - | $R_j$ | mRNA decay |
| - | $X_j$ | $l_j R_j$ | Translation |
| $X_j$ | - | $X_j$ | Protein decay |

Combining this positive feedback loop (PFL) module with the Chained-MMI2 Model yields a network in which genes 0 and 1 activate each other, gene 1 activates gene 2, and mRNAs 1 and 2 are each targeted by a microRNA at two sites. The gene index $j = 0, 1, 2$ is used for distinguishability with $i = 1, 2$, which applies only to microRNAs and their targets. MicroRNA-related reactions are as in the Chained-MMI2 Model. Only protein and mRNA dynamics are changed:

| Reactants | Products | Reaction Rate | Description |
|---|---|---|---|
| - | $R_0$ | $k_{R,0}\left(f_{0,C} + \frac{f_{0,1}\left(X_1/K_{0,1}\right)^{\mathrm{n}_{0,1}}}{1+\left(X_1/K_{0,1}\right)^{\mathrm{n}_{0,1}}}\right)$ | Regulated mRNA 0 transcription |
| - | $R_1$ | $k_{R,1}\left(f_{1,C} + \frac{f_{1,0}\left(X_0/K_{1,0}\right)^{\mathrm{n}_{1,0}}}{1+\left(X_0/K_{1,0}\right)^{\mathrm{n}_{1,0}}}\right)$ | Regulated mRNA 1 transcription |
| - | $R_2$ | $k_{R,2}\left(f_{2,C} + \frac{f_{2,1}\left(X_1/K_{2,1}\right)^{\mathrm{n}_{2,1}}}{1+\left(X_1/K_{2,1}\right)^{\mathrm{n}_{2,1}}}\right)$ | Regulated mRNA 2 transcription |
| $R_j$ | - | $R_j$ | mRNA decay |
| - | $X_j$ | $l_j R_j$ | Translation |
| $X_j$ | - | $X_j$ | Protein decay |
| MicroRNA-related reactions on mRNAs $i = 1, 2$ shared with the Chained-MMI2 Model | | | |

| Param. | Description | Figure 3B Red | | |
|---|---|---|---|---|
| | | Chain | PFL | Combined |
| $\alpha_{1,1}$ | RDF for mRNA 1 in the 1:1 complex with miRNA 1 | 2.580 | | 2.580 |
| $\alpha_{1,2}$ | RDF for mRNA 1 in the 1:2 complex | 9.112 | | 9.112 |
| $\alpha_{2,1}$ | RDF for mRNA 2 in the 1:1 complex with miRNA 2 | 2.235 | | 2.235 |
| $\alpha_{2,2}$ | RDF for mRNA 2 in the 1:2 complex | 7.544 | | 7.544 |
| $\beta_{1,1}$ | RDF for microRNA 1 in the 1:1 complex with mRNA 1 | 0.909 | | 0.909 |
| $\beta_{1,2}$ | RDF for microRNA 1 in the 1:2 complex | 0.326 | | 0.326 |
| $\beta_{2,1}$ | RDF for microRNA 2 in the 1:1 complex with mRNA 2 | 0.545 | | 0.545 |
| $\beta_{2,2}$ | RDF for microRNA 2 in the 1:2 complex | 0.227 | | 0.227 |
| $\gamma$ | Basal microRNA degradation rate | 1 | | 1 |
| $f_{0,C}$ | Proportion of mRNA 0 transcription rate allowed constitutively | | 0.028 | 0.028 |
| $f_{0,1}$ | Proportion of mRNA 0 transcription rate regulated by protein 1 | | 0.972 | 0.972 |
| $f_{1,C}$ | Proportion of mRNA 2 transcription rate allowed constitutively | | 0.097 | 0.097 |
| $f_{1,0}$ | Proportion of mRNA 2 transcription rate regulated by protein 0 | | 0.903 | 0.903 |
| $f_{2,C}$ | Proportion of mRNA 2 transcription rate allowed constitutively | 0.435 | | 0.435 |
| $f_{2,1}$ | Proportion of mRNA 2 transcription rate regulated by protein 1 | 0.565 | | 0.565 |
| $k_{r,1}$ | microRNA 1 transcription rate | 0.040 | | 0.040 |
| $k_{r,2}$ | microRNA 2 transcription rate | 0.109 | | 0.109 |
| $k_{R,0}$ | mRNA 0 maximum transcription rate | | 2.038 | 2.038 |
| $k_{R,1}$ | mRNA 1 maximum transcription rate | 0.389 | 1 | 0.920 |
| $k_{R,2}$ | mRNA 2 maximum transcription rate | 2.151 | | 2.151 |
| $K_1$ | Association constant between mRNA 1 and miRNA 1 | 5000 | | 5000 |
| $K_2$ | Association constant between mRNA 2 and miRNA 2 | 5000 | | 5000 |
| $K_{0,1}$ | Threshold of the regulation of mRNA 0 by protein 1 | | 3.738 | 3.738 |
| $K_{1,0}$ | Threshold of the regulation of mRNA 1 by protein 0 | | 0.144 | 0.144 |
| $K_{2,1}$ | Threshold of the regulation of mRNA 2 by protein 1 | 1.806 | | 1.806 |
| $l_0$ | Translation rate for mRNA 0 | | 1.463 | 1.463 |
| $l_1$ | Translation rate for mRNA 1 | 3.129 | 1.940 | 2.683 |
| $l_2$ | Translation rate for mRNA 2 | 3.423 | | 3.423 |
| $n_{0,1}$ | Cooperativity of regulation of mRNA 0 by protein 1 | | 3 | 3 |
| $n_{1,0}$ | Cooperativity of regulation of mRNA 1 by protein 0 | | 3 | 3 |
| $n_{2,1}$ | Cooperativity of regulation of mRNA 2 by protein 1 | 1 | | 1 |

#### MMI2 with transcriptional repression

This model includes one mRNA $R = C_0$, its protein product $X$, one microRNA $r$, and two mRNA-microRNA complexes $C_n$ for the number of microRNAs $n = 1, 2$. The protein represses the transcription of the microRNA.

| Reactants | Products | Reaction Rate | Description |
|---|---|---|---|
| - | $R$ | $k_R$ | mRNA transcription |
| $R$ | - | $R$ | mRNA decay |
| - | $X$ | $lR$ | Translation |
| $X$ | - | $X$ | Protein decay |
| - | $r$ | $k_r\left(f_{r,C} + \frac{f_{r,X}}{1+(X/K_{r,X})^{\mathrm{n}_{r,X}}}\right)$ | Regulated microRNA transcription then maturation |
| $r$ | - | $\gamma r$ | microRNA decay |
| $C_{n-1} + r$ | $C_n$ | $(3 - n)\kappa_{\mathrm{on}} C_{n-1} r$ | mRNA-microRNA binding |
| $C_n$ | $C_{n-1} + r$ | $n\kappa_{\mathrm{off}} C_n$ | mRNA-microRNA unbinding |
| $C_n$ | $nr$ | $\alpha_n C_n$ | Regulated mRNA degradation |
| $C_n$ | $C_{n-1}$ | $n\beta_n \gamma C_n$ | Regulated microRNA degradation |

| Parameter | Description | Figure 3D |
|---|---|---|
| $\alpha_1$ | RDF for mRNA in the 1:1 complex | 5 |
| $\alpha_2$ | RDF for mRNA in the 1:2 complex | 10 |
| $\beta_1$ | RDF for microRNA in the 1:1 complex | 0.65 |
| $\beta_2$ | RDF for microRNA in the 1:2 complex | 0.40 |
| $\gamma$ | Basal microRNA degradation rate | 1 |
| $f_{r,C}$ | Proportion of microRNA transcription rate allowed constitutively | 0.07 |
| $f_{r,X}$ | Proportion of microRNA transcription rate regulated by the protein | 0.93 |
| $k_r$ | microRNA maximum transcription rate | 0.07 |
| $k_R$ | mRNA transcription rate | Scanned |
| K | Association constant between mRNA and microRNA | 5000 |
| $K_{r.X}$ | Threshold of the regulation of microRNA by the protein | 0.9 |
| $l$ | mRNA translation rate | 2 |
| $\mathrm{n}_{r,X}$ | Cooperativity of the regulation of microRNA by the protein | 5 |

#### Two-microRNA MMI4 with transcriptional repressions

This model extends the two-microRNA MMI4 Model with a protein $X$ that is encoded by the mRNA and transcriptionally represses both microRNAs. Protein dynamics are added and microRNA transcription rates are changed, but other reactions are unaffected:

| **Reactants** | **Products** | **Reaction Rate** | **Description** |
|---|---|---|---|
| - | $X$ | $lR$ | Translation |
| $X$ | - | $X$ | Protein decay |
| - | $r_i$ | $k_{r,i}\left(f_{i,C} + \frac{f_{i,X}}{1+(X/K_{i,X})^{\mathrm{n}_{i,X}}}\right)$ | Regulated microRNA transcription then maturation |
| $r_i$ | - | $\gamma r_i$ | microRNA decay |
| Other mRNA and microRNA reactions shared with the two-microRNA MMI4 Model | | | |

| Parameter | Description | Fig. 3F Blue |
|---|---|---|
| $\alpha_{0,1}$ | RDF for mRNA in the 1:0:1 complex (one miRNA 2) | 4.164 |
| $\alpha_{0,2}$ | RDF for mRNA in the 1:0:2 complex | 4.559 |
| $\alpha_{1,0}$ | RDF for mRNA in the 1:1:0 complex | 1.125 |
| $\alpha_{1,1}$ | RDF for mRNA in the 1:1:1 complex | 5.483 |
| $\alpha_{1,2}$ | RDF for mRNA in the 1:1:2 complex | 13.577 |
| $\alpha_{2,0}$ | RDF for mRNA in the 1:2:0 complex | 2.250 |
| $\alpha_{2,1}$ | RDF for mRNA in the 1:2:1 complex | 4.646 |
| $\alpha_{2,2}$ | RDF for mRNA in the 1:2:2 complex | 5.981 |
| $\beta_{2,0,1}$ | RDF for miRNA 2 in the 1:0:1 complex | 0.482 |
| $\beta_{2,0,2}$ | RDF for miRNA 2 in the 1:0:2 complex | 0.231 |
| $\beta_{1,1,0}$ | RDF for miRNA 1 in the 1:1:0 complex | 0.325 |
| $\beta_{1,1,1}$ | RDF for miRNA 1 in the 1:1:1 complex | 0.190 |
| $\beta_{2,1,1}$ | RDF for miRNA 2 in the 1:1:1 complex | 0.531 |
| $\beta_{1,1,2}$ | RDF for miRNA 1 in the 1:1:2 complex | 0.714 |
| $\beta_{2,1,2}$ | RDF for miRNA 2 in the 1:1:2 complex | 0.109 |
| $\beta_{1,2,0}$ | RDF for miRNA 1 in the 1:2:0 complex | 0.061 |
| $\beta_{1,2,1}$ | RDF for miRNA 1 in the 1:2:1 complex | 0.070 |
| $\beta_{2,2,1}$ | RDF for miRNA 2 in the 1:2:1 complex | 0.795 |
| $\beta_{1,2,2}$ | RDF for miRNA 1 in the 1:2:2 complex | 0.041 |
| $\beta_{2,2,2}$ | RDF for miRNA 2 in the 1:2:2 complex | 0.167 |
| $\gamma$ | Basal microRNA degradation rate | 1 |
| $f_{1,C}$ | Proportion of miRNA 1 transcription rate allowed constitutively | 0.134 |
| $f_{1,X}$ | Proportion of miRNA transcription rate regulated by the protein | 0.866 |
| $f_{2,C}$ | Proportion of miRNA 2 transcription rate allowed constitutively | 0.059 |
| $f_{2,X}$ | Proportion of miRNA transcription rate regulated by the protein | 0.941 |
| $k_{r,1}$ | microRNA 1 maximum transcription rate | 0.108 |
| $k_{r,2}$ | microRNA 2 maximum transcription rate | 0.266 |
| $k_R$ | mRNA transcription rate | 2.776 |
| $K_1$ | Association constant between mRNA and microRNA 1 | 43810 |
| $K_2$ | Association constant between mRNA and microRNA 2 | 3906 |
| $K_{1.X}$ | Threshold of the regulation of microRNA 1 by the protein | 0.250 |
| $K_{2.X}$ | Threshold of the regulation of microRNA 2 by the protein | 5.684 |
| $l$ | mRNA translation rate | 7.916 |
| $n_{1,X}$ | Cooperativity of the regulation of microRNA 1 by the protein | 1 |
| $n_{2,X}$ | Cooperativity of the regulation of microRNA 2 by the protein | 4 |

#### Multi-feedback EMT network with MMI2

This model and the following are extensions of Subbalakshmi et al.'s EMT model [4], which uses a somewhat different form of equation for transcriptional regulation [5],

$$H^{-}(X,K,\mathrm{n})=\frac{1}{1+(X/K)^{\mathrm{n}}};$$

$$H^{s}(X,K,\mathrm{n},F)=H^{-}(X,K,\mathrm{n})+F\big(1-H^{-}(X,K,\mathrm{n})\big)$$

where $F$ is the fold-change in target expression resulting from overexpression of the regulator and the $H^{s}$ terms from multiple regulators are multiplied together.

This model includes *miR-200* microRNA $r_{200}$, *ZEB1* mRNA $R_{\mathrm{Z}}=C_{\mathrm{Z},0}$ encoding Zeb1, *SNAI1* mRNA $R_{\mathrm{Sn}}$ encoding Snail, *SNAI2* mRNA $R_{\mathrm{Sl}}$ encoding Slug, *CDH1* mRNA $R_{\mathrm{C}}$ encoding E-cadherin, the protein $X_{G}$ for each of the four protein-coding genes $G$, two complexes $C_{\mathrm{Z},n}$ for the number of *miR-200* molecules $n=1,2$ bound to a *ZEB1* mRNA, and a complex $C_{\mathrm{Sl}}$ of *miR-200* bound to *SNAI2* mRNA.

Cooperativities, fold changes, and protein degradation rates of transcription factors in the original Subbalakshmi et al. model were maintained. Fold changes of *CDH1* were as in the Chained-MMI2 Model.

| Reactants | Products | Reaction Rate | Description |
|---|---|---|---|
| - | $R_{\mathrm{C}}$ | $k_{\mathrm{C}} * H^s(X_{\mathrm{Z}}, K_{\mathrm{C,Z}}, n_{\mathrm{C,Z}}, F_{\mathrm{C,Z}}) * H^s(X_{\mathrm{Sn}}, K_{\mathrm{C,Sn}}, n_{\mathrm{C,Sn}}, F_{\mathrm{C,Sn}}) * H^s(X_{\mathrm{Sl}}, K_{\mathrm{C,Sl}}, n_{\mathrm{C,Sl}}, F_{\mathrm{C,Sl}})$ | Regulated *CDH1* transcription |
| - | $R_{\mathrm{Sl}}$ | $k_{\mathrm{Sl}} * H^s(X_{\mathrm{Sn}}, K_{\mathrm{Sl,Sn}}, n_{\mathrm{Sl,Sn}}, F_{\mathrm{Sl,Sn}}) * H^s(X_{\mathrm{Sl}}, K_{\mathrm{Sl,Sl}}, n_{\mathrm{Sl,Sl}}, F_{\mathrm{Sl,Sl}})$ | Regulated *SNAI2* transcription |
| - | $R_{\mathrm{Sn}}$ | $k_{\mathrm{Sn}} * H^s(X_{\mathrm{Sn}}, K_{\mathrm{Sn,Sn}}, n_{\mathrm{Sn,Sn}}, F_{\mathrm{Sn,Sn}}) * H^s(X_{\mathrm{Sl}}, K_{\mathrm{Sn,Sl}}, n_{\mathrm{Sn,Sl}}, F_{\mathrm{Sn,Sl}})$ | Regulated *SNAI1* transcription |
| - | $R_{\mathrm{Z}}$ | $k_{\mathrm{Z}} * H^s(X_{\mathrm{Z}}, K_{\mathrm{Z,Z}}, n_{\mathrm{Z,Z}}, F_{\mathrm{Z,Z}}) * H^s(X_{\mathrm{Sn}}, K_{\mathrm{Z,Sn}}, n_{\mathrm{Z,Sn}}, F_{\mathrm{Z,Sn}}) * H^s(X_{\mathrm{Sl}}, K_{\mathrm{Z,Sl}}, n_{\mathrm{Z,Sl}}, F_{\mathrm{Z,Sl}})$ | Regulated *ZEB1* transcription |
| $R_G$ | - | $R_G$ | mRNA decay |
| - | $X_{\mathrm{C}}$ | $l_{\mathrm{C}} R_{\mathrm{C}}$ | E-cadherin translation |
| - | $X_{\mathrm{Sl}}$ | $l_{\mathrm{Sl}}(R_{\mathrm{Sl}} + t_{\mathrm{Sl}} C_{\mathrm{Sl}})$ | Repressible Slug translation |
| - | $X_{\mathrm{Sn}}$ | $l_{\mathrm{Sn}} R_{\mathrm{Sn}}$ | Snail translation |
| - | $X_{\mathrm{Z}}$ | $l_{\mathrm{Z}}(R_{\mathrm{Z}} + t_{\mathrm{Z,1}} C_{\mathrm{Z,1}} + t_{\mathrm{Z,2}} C_{\mathrm{Z,2}})$ | Repressible Zeb1 translation |
| $X_G$ | - | $d_G X_G$ | Protein decay |
| - | $r_{200}$ | $k_{200} * H^s(X_{\mathrm{Z}}, K_{200,\mathrm{Z}}, n_{200,\mathrm{Z}}, F_{200,\mathrm{Z}}) * H^s(X_{\mathrm{Sn}}, K_{200,\mathrm{Sn}}, n_{200,\mathrm{Sn}}, F_{200,\mathrm{Sn}}) * H^s(X_{\mathrm{Sl}}, K_{200,\mathrm{Sl}}, n_{200,\mathrm{Sl}}, F_{200,\mathrm{Sl}})$ | Regulated *miR-200* transcription then maturation |
| $r_{200}$ | - | $\gamma r_{200}$ | *miR-200* decay |
| $R_{\mathrm{Sl}} + r_{200}$ | $C_{\mathrm{Sl}}$ | $\kappa_{\mathrm{on,Sl}} R_{\mathrm{Sl}} r_{200}$ | *SNAI2/miR-200* binding |
| $C_{\mathrm{Sl}}$ | $R_{\mathrm{Sl}} + r_{200}$ | $\kappa_{\mathrm{off,Sl}} C_{\mathrm{Sl}}$ | *SNAI2/miR-200* unbinding |
| $C_{\mathrm{Z},n-1} + r_{200}$ | $C_{\mathrm{Z},n}$ | $(3-n)\kappa_{\mathrm{on,Z}} C_{\mathrm{Z},n-1} r_{200}$ | *ZEB1/miR-200* binding |
| $C_{\mathrm{Z},n}$ | $C_{\mathrm{Z},n-1} + r_{200}$ | $n\kappa_{\mathrm{off,Z}} C_{\mathrm{Z},n}$ | *ZEB1/miR-200* unbinding |
| $C_{\mathrm{Sl}}$ | $r_{200}$ | $\alpha_{\mathrm{Sl}} C_{\mathrm{Sl}}$ | Regulated *SNAI2* decay |
| $C_{\mathrm{Sl}}$ | $R_{\mathrm{Sl}}$ | $\beta_{\mathrm{Sl}} \gamma C_{\mathrm{Sl}}$ | Regulated *miR-200* decay on *SNAI2* |
| $C_{\mathrm{Z},n}$ | $n r_{200}$ | $\alpha_{\mathrm{Z},n} C_{\mathrm{Z},n}$ | Regulated *ZEB1* decay |
| $C_{\mathrm{Z},n}$ | $C_{\mathrm{Z},n-1}$ | $n\beta_{\mathrm{Z},n} \gamma C_{\mathrm{Z},n}$ | Regulated *miR-200* decay on *ZEB1* |

| Parameter | Description | Figure 4C |
|---|---|---|
| $\alpha_{\mathrm{Sl}}$ | RDF for *SNAI2* in complex with *miR-200* | 8.890 |
| $\alpha_{\mathrm{Z,1}}$ | RDF for *ZEB1* in the 1:1 complex with *miR-200* | 5.048 |
| $\alpha_{\mathrm{Z,2}}$ | RDF for *ZEB1* in the 1:2 complex with *miR-200* | 12.431 |
| $\beta_{\mathrm{Sl}}$ | RDF for *miR-200* in complex with *SNAI2* | 0.093 |
| $\beta_{\mathrm{Z,1}}$ | RDF for *miR-200* in the 1:1 complex | 0.130 |
| $\beta_{\mathrm{Z,2}}$ | RDF for *miR-200* in the 1:2 complex | 0.101 |
| $\gamma$ | Basal microRNA degradation rate | 1 |
| $d_{\mathrm{C}}$ | Degradation rate of E-cadherin protein | 1 |
| $d_{\mathrm{Sl}}$ | Degradation rate of Slug protein | 1.10 |
| $d_{\mathrm{Sn}}$ | Degradation rate of Snail protein | 1.25 |
| $d_{\mathrm{Z}}$ | Degradation rate of Zeb1 protein | 1 |
| $F_{\mathrm{200,Sl}}$ | Fold-change in *miR-200* transcription due to Slug | 0.4 |
| $F_{\mathrm{200,Sn}}$ | Fold-change in *miR-200* transcription due to Snail | 0.1 |
| $F_{\mathrm{200,Z}}$ | Fold-change in *miR-200* transcription due to Zeb1 | 0.3 |
| $F_{\mathrm{C,Sl}}$ | Fold-change in *CDH1* transcription due to Slug | 0.3 |
| $F_{\mathrm{C,Sn}}$ | Fold-change in *CDH1* transcription due to Snail | 0.3 |
| $F_{\mathrm{C,Z}}$ | Fold-change in *CDH1* transcription due to Zeb1 | 0.75 |
| $F_{\mathrm{Sl,Sl}}$ | Fold-change in *SLUG* transcription due to Slug | 4 |
| $F_{\mathrm{Sl,Sn}}$ | Fold-change in *SLUG* transcription due to Snail | 0.5 |
| $F_{\mathrm{Sn,Sl}}$ | Fold-change in *SNAI* transcription due to Slug | 0.5 |
| $F_{\mathrm{Sn,Sn}}$ | Fold-change in *SNAI* transcription due to Snail | 4 |
| $F_{\mathrm{Z,Sl}}$ | Fold-change in *ZEB1* transcription due to Slug | 4 |
| $F_{\mathrm{Z,Sn}}$ | Fold-change in *ZEB1* transcription due to Snail | 10 |
| $F_{\mathrm{Z,Z}}$ | Fold-change in *ZEB1* transcription due to Zeb1 | 7.5 |
| $k_{200}$ | *miR-200* maximum transcription rate | 0.584 |
| $k_{\mathrm{C}}$ | *CDH1* basal transcription rate | 10 |
| $k_{\mathrm{Sl}}$ | *SNAI2* basal transcription rate | 0.383 |
| $k_{\mathrm{Sn}}$ | *SNAI1* basal transcription rate | 0.665 |
| $k_{\mathrm{Z}}$ | *ZEB1* basal transcription rate | 0.582 |
| $\mathrm{K_{Sl}}$ | Association constant between *SNAI2* and *miR-200* | 178 |
| $\mathrm{K_{Z}}$ | Association constant between *ZEB1* and *miR-200* | 2056 |
| $K_{\mathrm{200,Sl}}$ | Threshold of *miR-200* regulation by Slug | 13.034 |
| $K_{\mathrm{200,Sn}}$ | Threshold of *miR-200* regulation by Snail | 0.263 |
| $K_{\mathrm{200,Z}}$ | Threshold of *miR-200* regulation by Zeb1 | 0.127 |
| $K_{\mathrm{C,Sl}}$ | Threshold of *CDH1* regulation by Slug | 0.321 |
| $K_{\mathrm{C,Sn}}$ | Threshold of *CDH1* regulation by Snail | 0.147 |
| $K_{\mathrm{C,Z}}$ | Threshold of *CDH1* regulation by Zeb1 | 0.749 |
| $K_{\mathrm{Sl,Sl}}$ | Threshold of *SNAI2* regulation by Slug | 0.570 |
| $K_{\mathrm{Sl,Sn}}$ | Threshold of *SNAI2* regulation by Snail | 3.790 |
| $K_{\mathrm{Sn,Sl}}$ | Threshold of *SNAI1* regulation by Slug | 10.745 |
| $K_{\mathrm{Sn,Sn}}$ | Threshold of *SNAI1* regulation by Snail | 0.178 |

| | | |
|---|---|---|
| $K_{\mathrm{Z,Sl}}$ | Threshold of *ZEB1* regulation by Slug | 0.046 |
| $K_{\mathrm{Z,Sn}}$ | Threshold of *ZEB1* regulation by Snail | 0.806 |
| $K_{\mathrm{Z,Z}}$ | Threshold of *ZEB1* regulation by Zeb1 | 7.792 |
| $l_{\mathrm{C}}$ | *CDH1* translation rate | 1 |
| $l_{\mathrm{Sl}}$ | Free *SNAI2* translation rate | 0.746 |
| $l_{\mathrm{Sn}}$ | *SNAI1* translation rate | 0.331 |
| $l_{\mathrm{Z}}$ | Free *ZEB1* translation rate | 0.737 |
| $\mathrm{n}_{200,\mathrm{Sl}}$ | Cooperativity of *miR-200* regulation by Slug | 1 |
| $\mathrm{n}_{200,\mathrm{Sn}}$ | Cooperativity of *miR-200* regulation by Snail | 2 |
| $\mathrm{n}_{200,\mathrm{Z}}$ | Cooperativity of *miR-200* regulation by Zeb1 | 3 |
| $\mathrm{n}_{\mathrm{C,Sl}}$ | Cooperativity of *CDH1* regulation by Slug | 2 |
| $\mathrm{n}_{\mathrm{C,Sn}}$ | Cooperativity of *CDH1* regulation by Snail | 2 |
| $\mathrm{n}_{\mathrm{C,Z}}$ | Cooperativity of *CDH1* regulation by Zeb1 | 2 |
| $\mathrm{n}_{\mathrm{Sl,Sl}}$ | Cooperativity of *SNAI2* regulation by Slug | 4 |
| $\mathrm{n}_{\mathrm{Sl,Sn}}$ | Cooperativity of *SNAI2* regulation by Snail | 1 |
| $\mathrm{n}_{\mathrm{Sn,Sl}}$ | Cooperativity of *SNAI1* regulation by Slug | 3 |
| $\mathrm{n}_{\mathrm{Sn,Sn}}$ | Cooperativity of *SNAI1* regulation by Snail | 5 |
| $\mathrm{n}_{\mathrm{Z,Sl}}$ | Cooperativity of *ZEB1* regulation by Slug | 2 |
| $\mathrm{n}_{\mathrm{Z,Sn}}$ | Cooperativity of *ZEB1* regulation by Snail | 2 |
| $\mathrm{n}_{\mathrm{Z,Z}}$ | Cooperativity of *ZEB1* regulation by Zeb1 | 2 |
| $t_{\mathrm{Sl}}$ | Fold-change in *SNAI2* translation in complex with *miR-200* | 0.146 |
| $t_{\mathrm{Z,1}}$ | Fold-change in *ZEB1* translation in 1:1 complex with *miR-200* | 0.301 |
| $t_{\mathrm{Z,2}}$ | Fold-change in *ZEB1* translation in 1:2 complex with *miR-200* | 0.067 |

### Multi-feedback EMT network with MMI2 and miR-101

This model extends the previous by adding *miR-101* microRNA $r_{101}$ which targets *ZEB1* at one binding site to form complexes $C_{\mathrm{Z},m,n}$ where $m$ is the number of bound *miR-101* molecules and $n = 0, 1$ is the number of bound *miR-200* molecules. Slug and Snail proteins repress *miR-101*. Only *miR-101* and *ZEB1*-microRNA-related dynamics are added/changed:

| Reactants | Products | Reaction Rate | Description |
|---|---|---|---|
| - | $X_{\mathrm{Z}}$ | $l_{\mathrm{Z}}\left(R_{\mathrm{Z}} + \sum_{(m,n)\neq(0,0)} t_{\mathrm{Z},m,n} C_{\mathrm{Z},m,n}\right)$ | Repressible Zeb1 translation |
| - | $r_{101}$ | $k_{101} *$ $H^{s}\left(X_{\mathrm{Sn}}, K_{101,\mathrm{Sn}}, n_{101,\mathrm{Sn}}, F_{101,\mathrm{Sn}}\right) *$ $H^{s}\left(X_{\mathrm{Sl}}, K_{101,\mathrm{Sl}}, n_{101,\mathrm{Sl}}, F_{101,\mathrm{Sl}}\right)$ | Regulated *miR-101* transcription then maturation |
| $r_{101}$ | - | $\gamma r_{101}$ | *miR-101* decay |
| $C_{\mathrm{Z},0,n} + r_{101}$ | $C_{\mathrm{Z},1,n}$ | $\kappa_{\mathrm{on,Z},101} C_{\mathrm{Z},0,n} r_{101}$ | *ZEB1/miR-101* binding |
| $C_{\mathrm{Z},1,n}$ | $C_{\mathrm{Z},0,n} + r_{101}$ | $\kappa_{\mathrm{off,Z},101} C_{\mathrm{Z},1,n}$ | *ZEB1/miR-101* unbinding |
| $C_{\mathrm{Z},m,n-1} + r_{200}$ | $C_{\mathrm{Z},m,n}$ | $(3-n)\kappa_{\mathrm{on,Z},200} C_{\mathrm{Z},m,n-1} r_{200}$ | *ZEB1/miR-200* binding |
| $C_{\mathrm{Z},m,n}$ | $C_{\mathrm{Z},m,n-1} + r_{200}$ | $n\kappa_{\mathrm{off,Z},200} C_{\mathrm{Z},m,n}$ | *ZEB1/miR-200* unbinding |
| $C_{\mathrm{Z},m,n}$ | $m r_{101} + n r_{200}$ | $\alpha_{\mathrm{Z},m,n} C_{\mathrm{Z},m,n}$ | Regulated *ZEB1* decay |
| $C_{\mathrm{Z},1,n}$ | $C_{\mathrm{Z},0,n}$ | $n\beta_{\mathrm{Z},101,1,n} \gamma C_{\mathrm{Z},1,n}$ | Regulated *miR-101* decay on *ZEB1* |
| $C_{\mathrm{Z},m,n}$ | $C_{\mathrm{Z},m,n-1}$ | $n\beta_{\mathrm{Z},200,m,n} \gamma C_{\mathrm{Z},m,n}$ | Regulated *miR-200* decay on *ZEB1* |
| Dynamics of other mRNAs, proteins, and *miR-200* interactions shared with previous model | | | |

| Parameter | Description | Fig. 4F | Fig. 4G |
|---|---|---|---|
| $\alpha_{\mathrm{Sl}}$ | RDF for *SNAI2* in complex with *miR-200* | 1.732 | 1.788 |
| $\alpha_{\mathrm{Z},0,1}$ | RDF for *ZEB1* in the 1:0:1 complex (one *miR-200*) | 2.080 | 1.037 |
| $\alpha_{\mathrm{Z},0,2}$ | RDF for *ZEB1* in the 1:0:2 complex | 2.965 | 2.864 |
| $\alpha_{\mathrm{Z},1,0}$ | RDF for *ZEB1* in the 1:1:0 complex (one *miR-101*) | 3.149 | 6.022 |
| $\alpha_{\mathrm{Z},1,1}$ | RDF for *ZEB1* in the 1:1:1 complex | 5.330 | 12.066 |
| $\alpha_{\mathrm{Z},1,2}$ | RDF for *ZEB1* in the 1:1:2 complex | 9.704 | 12.066 |
| $\beta_{\mathrm{Sl}}$ | RDF for *miR-200* in complex with *SNAI2* | 0.192 | 0.154 |
| $\beta_{\mathrm{Z},200,0,1}$ | RDF for *miR-200* in the 1:0:1 complex with *ZEB1* | 0.607 | 0.174 |
| $\beta_{\mathrm{Z},200,0,2}$ | RDF for *miR-200* in the 1:0:2 complex with *ZEB1* | 0.260 | 0.224 |
| $\beta_{\mathrm{Z},101,1,0}$ | RDF for *miR-101* in the 1:1:0 complex with *ZEB1* | 0.326 | 0.602 |
| $\beta_{\mathrm{Z},101,1,1}$ | RDF for *miR-101* in the 1:1:1 complex with *ZEB1* | 0.119 | 0.478 |
| $\beta_{\mathrm{Z},200,1,1}$ | RDF for *miR-200* in the 1:1:1 complex with *ZEB1* | 0.294 | 0.065 |
| $\beta_{\mathrm{Z},101,1,2}$ | RDF for *miR-101* in the 1:1:2 complex with *ZEB1* | 0.126 | 0.440 |
| $\beta_{\mathrm{Z},200,1,2}$ | RDF for *miR-200* in the 1:1:2 complex with *ZEB1* | 0.101 | 0.134 |
| $\gamma$ | Basal microRNA degradation rate | 1 | 1 |
| $d_{\mathrm{C}}$ | Degradation rate of E-cadherin protein | 1 | 1 |
| $d_{\mathrm{Sl}}$ | Degradation rate of Slug protein | 1.10 | 1.10 |
| $d_{\mathrm{Sn}}$ | Degradation rate of Snail protein | 1.25 | 1.25 |
| $d_{\mathrm{Z}}$ | Degradation rate of Zeb1 protein | 1 | 1 |
| $F_{101,\mathrm{Sl}}$ | Fold-change in *miR-101* transcription due to Slug | 0.4 | 0.4 |
| $F_{101,\mathrm{Sn}}$ | Fold-change in *miR-101* transcription due to Snail | 0.1 | 0.1 |
| $F_{200,\mathrm{Sl}}$ | Fold-change in *miR-200* transcription due to Slug | 0.4 | 0.4 |
| $F_{200,\mathrm{Sn}}$ | Fold-change in *miR-200* transcription due to Snail | 0.1 | 0.1 |
| $F_{200,\mathrm{Z}}$ | Fold-change in *miR-200* transcription due to Zeb1 | 0.1 | 0.1 |
| $F_{\mathrm{C,Sl}}$ | Fold-change in *CDH1* transcription due to Slug | 0.3 | 0.3 |
| $F_{\mathrm{C,Sn}}$ | Fold-change in *CDH1* transcription due to Snail | 0.3 | 0.3 |
| $F_{\mathrm{C,Z}}$ | Fold-change in *CDH1* transcription due to Zeb1 | 0.75 | 0.75 |
| $F_{\mathrm{Sl,Sl}}$ | Fold-change in *SNAI2* transcription due to Slug | 4 | 4 |
| $F_{\mathrm{Sl,Sn}}$ | Fold-change in *SNAI2* transcription due to Snail | 0.5 | 0.5 |
| $F_{\mathrm{Sn,Sl}}$ | Fold-change in *SNAI1* transcription due to Slug | 0.5 | 0.5 |
| $F_{\mathrm{Sn,Sn}}$ | Fold-change in *SNAI1* transcription due to Snail | 0.4 | 0.4 |
| $F_{\mathrm{Z,Sl}}$ | Fold-change in *ZEB1* transcription due to Slug | 4 | 4 |
| $F_{\mathrm{Z,Sn}}$ | Fold-change in *ZEB1* transcription due to Snail | 10 | 10 |
| $F_{\mathrm{Z,Z}}$ | Fold-change in *ZEB1* transcription due to Zeb1 | 7.5 | 7.5 |
| $k_{101}$ | *miR-101* basal transcription rate | 0.981 | 0.859 |
| $k_{200}$ | *miR-200* basal transcription rate | 0.584 | 1.531 |
| $k_{\mathrm{C}}$ | *CDH1* basal transcription rate | 10 | 10 |
| $k_{\mathrm{Sl}}$ | *SNAI2* basal transcription rate | 0.444 | 0.877 |
| $k_{\mathrm{Sn}}$ | *SNAI1* basal transcription rate | 1.593 | 1.090 |
| $k_{\mathrm{Z}}$ | *ZEB1* basal transcription rate | 1.741 | 0.338 |
| $\mathrm{K}_{\mathrm{Sl}}$ | Association constant between *SNAI2* and *miR-200* | 5965 | 126 |

| | | | |
|---|---|---|---|
| $K_{Z,101}$ | Association constant between *ZEB1* and *miR-101* | 69231 | 455 |
| $K_{Z,200}$ | Association constant between *ZEB1* and *miR-200* | 83360 | 1009 |
| $K_{101,Sl}$ | Threshold of *miR-101* regulation by Slug | 3.355 | 7.175 |
| $K_{101,Sn}$ | Threshold of *miR-101* regulation by Snail | 7.782 | 0.187 |
| $K_{200,Sl}$ | Threshold of *miR-200* regulation by Slug | 0.173 | 10.586 |
| $K_{200,Sn}$ | Threshold of *miR-200* regulation by Snail | 57.659 | 55.047 |
| $K_{200,Z}$ | Threshold of *miR-200* regulation by Zeb1 | 2.767 | 22.856 |
| $K_{C,Sl}$ | Threshold of *CDH1* regulation by Slug | 0.623 | 0.317 |
| $K_{C,Sn}$ | Threshold of *CDH1* regulation by Snail | 1.291 | 0.719 |
| $K_{C,Z}$ | Threshold of *CDH1* regulation by Zeb1 | 2.957 | 0.034 |
| $K_{Sl,Sl}$ | Threshold of *SNAI2* regulation by Slug | 1.198 | 0.254 |
| $K_{Sl,Sn}$ | Threshold of *SNAI2* regulation by Snail | 52.715 | 0.213 |
| $K_{Sn,Sl}$ | Threshold of *SNAI1* regulation by Slug | 3.043 | 38.719 |
| $K_{Sn,Sn}$ | Threshold of *SNAI1* regulation by Snail | 6.441 | 34.999 |
| $K_{Z,Sl}$ | Threshold of *ZEB1* regulation by Slug | 0.253 | 6.644 |
| $K_{Z,Sn}$ | Threshold of *ZEB1* regulation by Snail | 4.445 | 0.987 |
| $K_{Z,Z}$ | Threshold of *ZEB1* regulation by Zeb1 | 81.449 | 0.037 |
| $l_C$ | *CDH1* translation rate | 1 | 1 |
| $l_{Sl}$ | Free *SNAI2* translation rate | 1.298 | 2.388 |
| $l_{Sn}$ | *SNAI1* translation rate | 1.017 | 0.825 |
| $l_Z$ | Free *ZEB1* translation rate | 2.597 | 0.726 |
| $n_{101,Sl}$ | Cooperativity of *miR-101* regulation by Slug | 1 | 1 |
| $n_{101,Sn}$ | Cooperativity of *miR-101* regulation by Snail | 2 | 2 |
| $n_{200,Sl}$ | Cooperativity of *miR-200* regulation by Slug | 1 | 1 |
| $n_{200,Sn}$ | Cooperativity of *miR-200* regulation by Snail | 2 | 2 |
| $n_{200,Z}$ | Cooperativity of *miR-200* regulation by Zeb1 | 3 | 3 |
| $n_{C,Sl}$ | Cooperativity of *CDH1* regulation by Slug | 2 | 2 |
| $n_{C,Sn}$ | Cooperativity of *CDH1* regulation by Snail | 2 | 2 |
| $n_{C,Z}$ | Cooperativity of *CDH1* regulation by Zeb1 | 2 | 2 |
| $n_{Sl,Sl}$ | Cooperativity of *SNAI2* regulation by Slug | 4 | 4 |
| $n_{Sl,Sn}$ | Cooperativity of *SNAI2* regulation by Snail | 1 | 1 |
| $n_{Sn,Sl}$ | Cooperativity of *SNAI1* regulation by Slug | 3 | 3 |
| $n_{Sn,Sn}$ | Cooperativity of *SNAI1* regulation by Snail | 5 | 5 |
| $n_{Z,Sl}$ | Cooperativity of *ZEB1* regulation by Slug | 2 | 2 |
| $n_{Z,Sn}$ | Cooperativity of *ZEB1* regulation by Snail | 2 | 2 |
| $n_{Z,Z}$ | Cooperativity of *ZEB1* regulation by Zeb1 | 2 | 2 |
| $t_{Sl}$ | Fold-change in *SNAI2* translation in complex with *miR-200* | 0.914 | 0.150 |
| $t_{Z,0,1}$ | Fold-change in *ZEB1* translation in the 1:0:1 complex | 0.079 | 0.166 |
| $t_{Z,0,2}$ | Fold-change in *ZEB1* translation in the 1:0:2 complex | 0.050 | 0.047 |
| $t_{Z,1,0}$ | Fold-change in *ZEB1* translation in the 1:1:0 complex | 0.061 | 0.564 |
| $t_{Z,1,1}$ | Fold-change in *ZEB1* translation in the 1:1:1 complex | 0.013 | 0.093 |
| $t_{Z,1,2}$ | Fold-change in *ZEB1* translation in the 1:1:2 complex | 0.0002 | 0.047 |
